# Time-Resolved Crystallography Reveals the Mechanisms of GTP hydrolysis for N-RAS and the Oncogenic Mutants G12C, G12V and Q61L

**DOI:** 10.1101/2025.08.21.670574

**Authors:** Guowu Lin, Paola Zinser-Peniche, Xiaohong Zhou, Ulises Santiago, Igor Kurnikov, Silvia Russi, Brisa Chagas, May E. Sharpe, Dennis P. Stegmann, Shannon A. Heinig, Rohan K. Goyal, V. Nagarajan, Daniel Calero, Sandra Vergara, Alexander Deiters, Maria G. Kurnikova, Timothy F. Burns, Aina Cohen, Guillermo Calero

## Abstract

The RAS family of small GTPases are molecular switches that convey downstream signals regulating cell proliferation, differentiation, and apoptosis. The signaling competent GTP-bound RAS transitions to its inactive GDP-bound form through γ-phosphate hydrolysis. Oncogenic RAS mutations hamper GTP hydrolysis and are present in up to 30% of all human cancers. Structural studies of RAS proteins bound to non-hydrolysable GTP analogs have revealed snapshots of the enzyme in its possibly active form. Yet, the mechanism of GTP hydrolysis has not been structurally resolved. To visualize this reaction in real time, we performed time-resolved crystallographic experiments employing a photolabile caged-GTP substrate. Fifty-seven distinctive reaction intermediates were captured during hydrolysis of a live GTP for N-RAS, the oncogenic mutants G12C, G12V and Q61L; and Y32R, a fast hydrolytic mutant. The reaction mechanisms and rates for the native and each of the mutants differed significantly; however, they shared common elements: an initially catalytically-defective open state, which transitions into the closed Michaelis complex state with solvent-assisted O3B-Pγ bond lengthening and breaking, followed by the release of the Mg^2+^ stabilized PO3^-^/PO4^-3^ species and unfolding of the switch loops. Given the conserved nature of GTP- and ATP-ases active sites, this structural work lays the basis to understand the universal mechanism of γ-phosphate hydrolysis. Furthermore, search for cryptic binding sites during GTP hydrolysis in G12C, G12V, and Q61L mutants reveals the presence of distinctive state-dependent binding pockets that could be targets for structure-based drug discovery of experimentally resolved intermediates states.

---

The monomeric GTPases are a ubiquitous family of enzymes that catalyze hydrolysis of guanosine triphosphate (GTP) into guanosine diphosphate (GDP)^1^. The active, GTP-bound form of the RAS family of GTPases, including the three human RAS genes K-, H- and N-RAS, initiate signaling pathways that trigger cellular proliferation. RAS mutations that interfere with GTP hydrolysis are often observed in human malignancies^2–7^. GTP hydrolysis can be intrinsic, which is typically slow (0.0068s^-1^)^5,8^, or RAS-GAP mediated, which occurs two orders of magnitude faster (0.8-5 s^-1^)^9–13^. The mechanism of RAS-GAP-mediated hydrolysis was illuminated by the crystal structure of the RAS-GAP bound to the K-RAS-GDP-AlF_3_ complex^14^, where RAS-GAP residues position K-RAS Gln^61^ and RAS-GAP residue Arg^789^ next to AlF3 to trigger hydrolysis. Despite the wealth of structural information for the RAS^15,16^ (and all other) families of GTPases, observation of the mechanism of intrinsic GTP hydrolysis has remained elusive. Earlier structural work has been limited by the necessary use of non-hydrolysable GTP analogues and by crystal contacts that restrain or position the catalytic loops, switch 1 (SW1, residues 30-38) and switch 2 (SW2, residues 59-72), in non-productive conformations; hence, the architecture of the RAS-active site in the presence of a “live” GTP and the conformational changes that trigger GTP hydrolysis have resisted observation^17^. Other outstanding questions involve the position of SW1 and SW2 loops in the GTP- and GDP-bound states, their conformational changes during catalysis, and the nature of the GTP hydrolysis mechanism. Two possible limiting cases for the reaction transition state have been postulated for GTP hydrolysis: a dissociative and an associative^18,19^. In the former, the leaving group departure precedes nucleophilic attack, forming a trigonal metaphosphate (PO^3-^); in the latter, the catalytic water (W_cat_) performs a nucleophilic attack on the γ-phosphate (Pγ) atom of GTP before the elongation or breaking of Pγ–O3B bond, forming a penta-coordinated phosphorous transition state^20,21^. FTIR studies using caged substrates and QM/MM simulations of GTP-bound K-RAS have shown that the dissociative reaction is energetically favored^9,22–24^; however, no structural evidence for either state is available. Direct observation of the overall conformation of the switches, the active site architecture, the mechanisms of GTP hydrolysis, and the effect of oncogenic mutations to these processes is essential to drive the development of new therapies for RAS-based malignancies and developmental defects^25^.

Here we use time-resolved crystallographic studies to create a structural movie of GTP hydrolysis using the small GTPase N-RAS and UV-photolysis of photoactive NPE-caged GTP. The movie reveals a long-lived, post-photolysis reactant state with an open active site pocket and reduced hydrolysis rate. A hinged motion of SW1 residues closes the active site positioning Tyr^32^ (SW1) next to the γ-phosphate and allowing Gln^61^ (SW2) to steer the catalytic water for γ-phosphate nucleophilic attack. Intriguingly, a PO3^-^/PO4^-33^ species was captured during the reaction and was stabilized through Mg^2+^ coordination and H-bonds with P-loop, SW1 and SW2 residues. Release of PO3^-^/PO4^-3^ species triggers SW1 and SW2 unfolding in the GDP product state. Crystallization conditions with freely moving SW1 and SW2 residues were reproduced for P-loop G12C and G12V, and SW2 Q61L mutants, elucidating the structural basis for their diminished catalytic activities. The structural movies allowed development of quantitative models of GTP hydrolysis, including mechanism and rates, using extensive free energy molecular dynamics (MD) simulations with the empirical valence bond (EVB) model for N-RAS and mutants^11,26,27^. Discovery of a mutant with high hydrolytic activity (135-fold faster) Y32R showed low proliferation rates and prevented NRAS G12C mutant driven transformation.

## Caged GTP photolysis observed in crystals exposed to 365 nm light

Previous studies of H-RAS bound to the photolabile P^3^-(1-(2-nitrophenyl)-ethyl)-ester-caged guanosine-5’-triphosphate (NPE-caged GTP) followed by photolysis allowed for observation of the first “live” GTP in the H-RAS active site; however, crystal contacts limited SW1 mobility hampering observation of the events leading to hydrolysis^28^. To overcome this limitation, we postulated that crystallization conditions with unrestrained switch 1 (SW1) and switch 2 (SW2) mobility, could be conducive to observation of GTP hydrolysis. Moreover, low ionic strength, room temperature crystallization (since the reaction is intrinsically slow^29^) and no phosphate ions (that could introduce bias) were highly desirable. We identified crystallization conditions for K-RAS and N-RAS bound to NPE-caged GTP substrate. Crystals were UV-photolyzed (at 365 nm) to remove the NPE-cage and frozen at varied time points (**Extended Data Fig. 1a**). The difference map (F_obs_ – F_calc_) from an initial K-RAS control or caged data set (before UV-photolysis) showed electron density for NPE-caged GTP with the nitrobenzyl group involved in packing interactions with P-loop residues (**Extended Data Fig. 1b**). Data sets collected following UV exposure revealed efficient NPE photolysis, however, no GTP hydrolysis was observed for K-RAS since Tyr^32^ was trapped inside a small pocket formed by symmetry related molecules that limited SW1 mobility; and Thr^35^ did not form part of the Mg^2+^ coordination sphere (**Extended Data Fig. 1c,d**). These experiments highlighted the flexibility of SW1 and SW2 residues and corroborated the critical role of Tyr^32^ and Thr^35^ in the hydrolytic process^30^.

## Time-resolved studies reveal the mechanisms of GTP hydrolysis by N-RAS

We found that crystals of NPE-caged GTP bound N-RAS had no symmetry-related contacts involving the switch loops allowing free mobility (**Extended Data Fig. 1e**); the overall B-factors of the switch loops were on average higher than the rest of the structure indicating enhanced flexibility of SW1 and SW2 (**Extended Data Fig. 1f**). Several dozen N-RAS-WT data sets were collected, but we limited our analysis to those crystals diffracting between 1.6 to 1.9 Å resolution. Examination of the electron density enabled classification of the structural data under three criteria: the conformation of SW1 and SW2 residues (**Fig. 1a**); the extent of the electron density around the oxygen atom (O3B) bonding the β-γ phosphates (**Fig. 1b**); and the occupancy of the γ-phosphate (**Fig. 1c**). Six major states were identified (**Extended Data Fig. 1g**): 1) The NPE-GTP bound (pre-photolysis) control or “caged” state (**Extended Data Fig. 1h**); 2) the reactant post-photolysis state (**Fig. 1a, I-IV**) with an open active site and solvent exposed GTP (hereafter, open state); 3) the Michaelis complex state (hereafter, closed state) resulting from the hinged motion of SW1 residues closing the active site (**Fig. 1a, V-VII**); 4) the hydrolysis state with O3B-γ-phosphate bond lengthening and breaking (**Fig. 1a, VIII-IX**); 5) the PO3^-^/PO4^3-^ release state (**Fig. 1a, X-XII**) with a trapped PO4^3-^ ion (**Fig. 1a, XIII**); and 6) the GDP “product state” with unfolded SW1 and SW2 loops (**Fig. 1a, XIV**). The time for GTP hydrolysis, from photolysis of the caged compound to product release in the crystals (at 16 °C), was approximately 18 hours, which is consistent with experimental results at lower hydrolysis temperatures^31^. In total, fifteen data sets belonging to the six distinct major states were captured in these experiments.

**Fig. 1:**
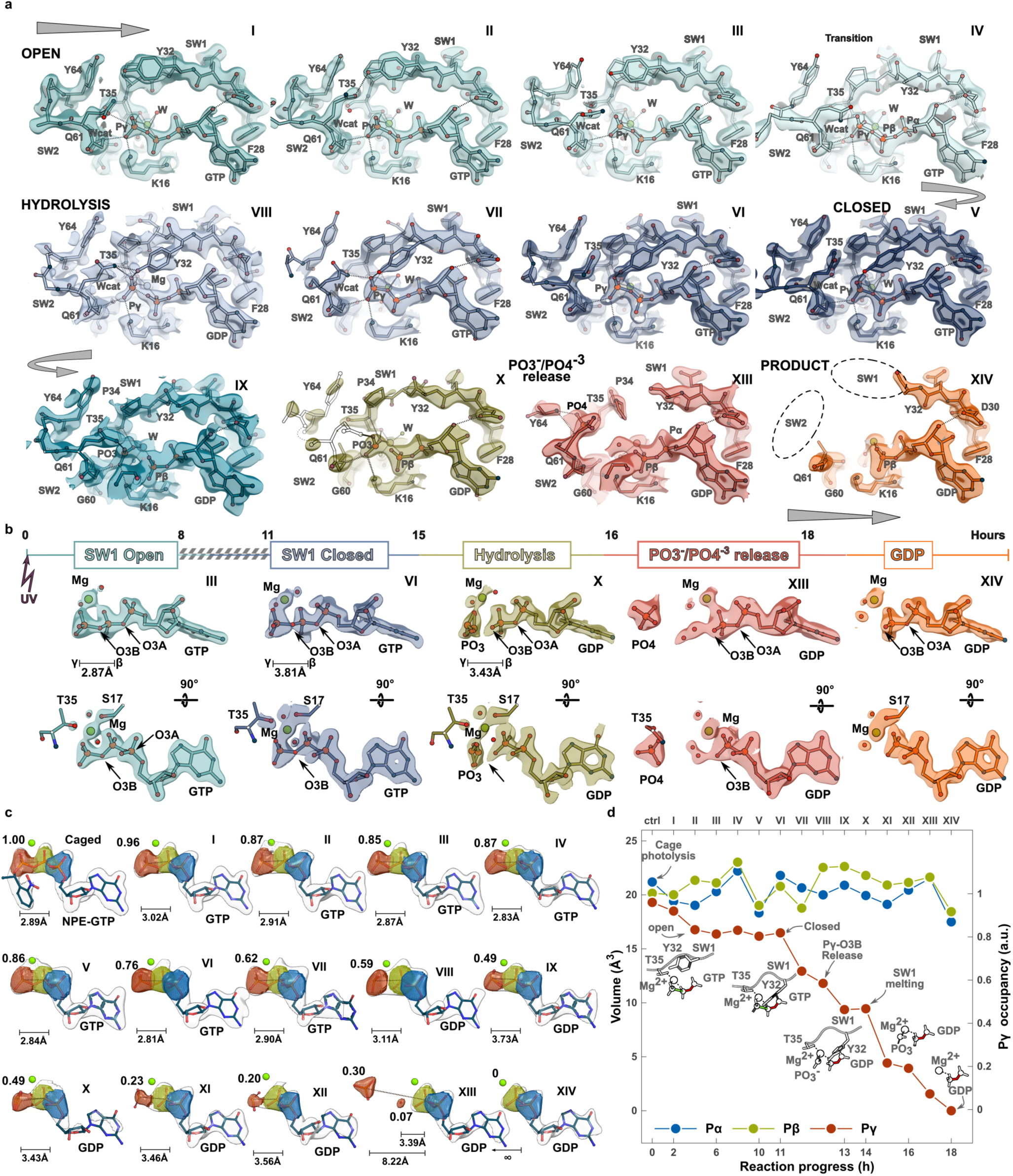
Structural classification of GTP hydrolysis. **a**, *<u>Switch view</u>*. 2Fo-Fc maps contoured at 1.5 0 for post-photolysis structures illustrating the N-RAS active site with SW1 and SW2 in open (cyan), closed (blue) or disordered conformation (green-orange). **b,** Two views of the 2Fo-Fc maps contoured at 2.0 0 around GTP and B-factor sharpening of −10; arrows point to density around O3A and O3B atoms for comparison. **c**, Polder (difference) maps contoured at 4.5 0 omitting GTP from the calculation. Blue, green and red spheres correspond to α- β- and γ-phosphates, respectively. Bar illustrates the β-γ bond distance. **d**, Occupancy of the γ-phosphate as function of the reaction process.

The “caged state” (NPE GTP-bound) shows well-defined electron density for SW1, SW2 and the NPE-caged GTP (**Extended Data Fig. 1h**). The NPE moiety is sandwiched between Tyr^32^ and Gly^13^, and forms H-bonds with the amide backbone and the carbonyl of Asp^33^ (**Extended Data Fig. 1i**). SW1 residue Thr^35^ forms part of the octahedral Mg^2+^ coordination, as commonly observed in previous RAS crystal structures bound to non-hydrolysable GTP analogs^32^ (**Extended Data Fig. 1j,k**). The catalytic water (W_cat_) was captured in the caged state and was found approximately 3.6 Å away from the γ-phosphate (**Extended Data Fig. 1i**).

The open state reveals a solvent exposed GTP in the active site; however, no major changes with respect to the NPE-GTP structure are seen (**Extended Data Fig. 2a**). The open conformation is long-lived since it is seen up to 11 hours post-photolysis (**Extended Data Fig. 1g**). The structures show small positional changes in SW1 and SW2 residues, as well as minor changes in bond lengths and α-, β- and γ-phosphate dihedral angles (**Fig. 2a**); Mg^2+^ coordination by the β- and γ-phosphates is bidentate with the O2B-Mg^2+^ distance slightly shorter than O2G-Mg^2+^ (**Extended Data Fig. 2b**); however, nearing the end of the open-state, the β-γ phosphate angle narrows and the O2G-Mg^2+^ distance shortens while the O2B-Mg^2+^ distance lengthens (**Extended Data Fig. 2c,d**). These changes in Mg^2+^ coordination distance could lead to a net charge transfer from the γ-phosphate O2G to Mg^2+^, weakening the Pγ-03Β bond^24^. Polder difference map analysis (**Fig. 1d**) shows that the overall occupancy of the γ-phosphate (compared to the α- and β-phosphates) during the open state drops to around 85%, indicating that a small fraction of the N-RAS molecule population within the crystal has undergone hydrolysis.

**Fig. 2:**
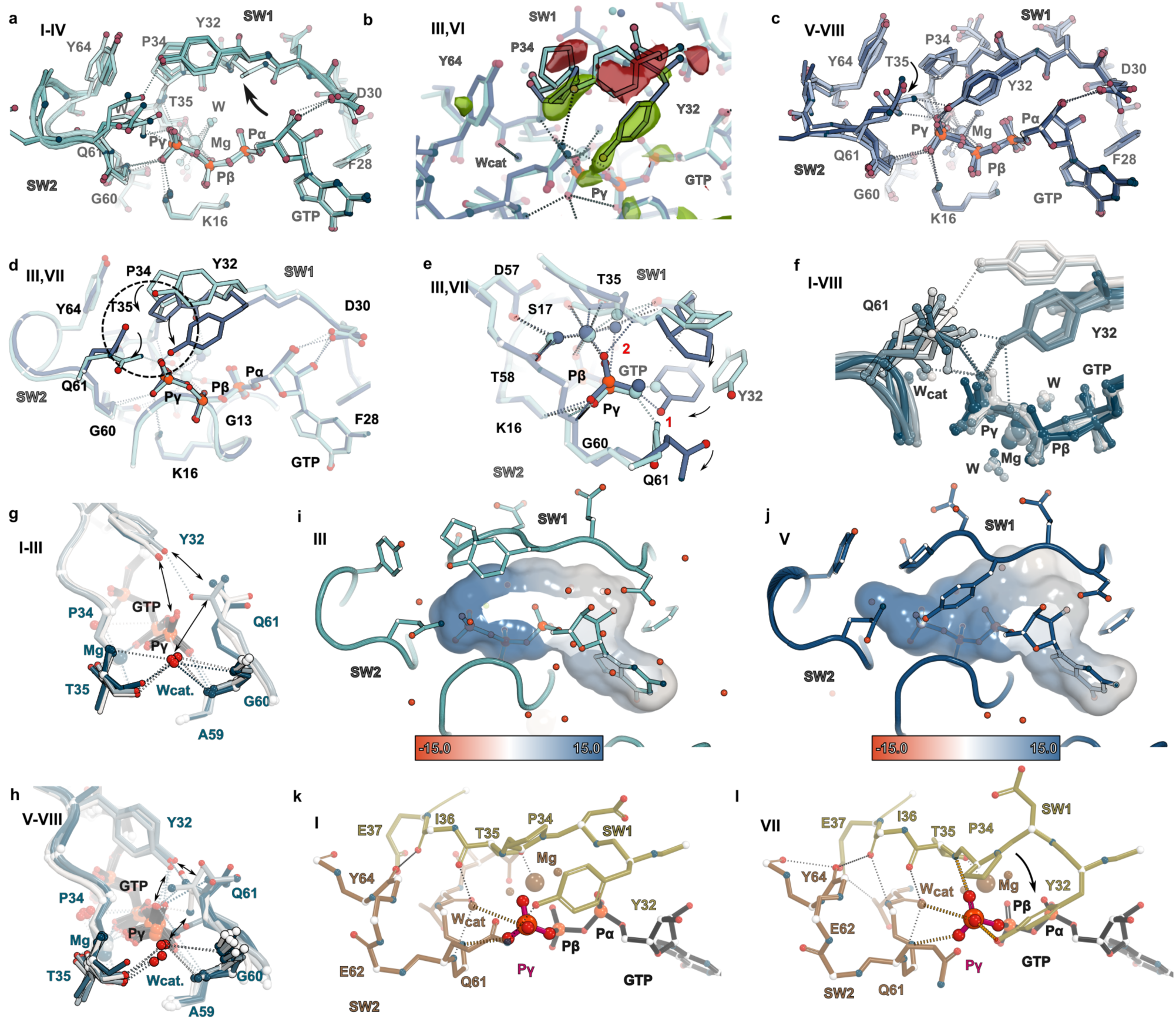
Conformational changes during open to closed transition. **a**, *<u>Switch view</u>*, cartoon and ball and stick representation of the overlay between four open state structures. Dashed lines hereafter indicate hydrogen bonds with distances below 3.5 Å. **b**, Difference maps contoured at +/- 30. Negative (red) and positive (green) density around SW1 residues Tyr^32^-Pro^34^ denote the structural changes seen during the open-to-closed transition. **c**, Cartoon and ball and stick representation of the overlay between four closed state structures. **d**, *<u>switch-</u>* and **e**, *<u>γ-</u>*phosphate-views of the overlay between the open (state III, cyan) and closed (state VII, blue) structures depicting the conformational changes occurring during open-to-close transition; dashed circle highlights critical residues, red numbers indicate H-bonds formed in the closed state). **f**, Rotamers of Gln^61^ in open and closed states (light to blue colors), illustrating its high mobility. **g**,**h**, *<u>γ-phosphate view</u>* of the W_cat_ stabilization by neighboring residues in the open (**g**) and closed (**h**) conformations. **i**,**j**, Surface representation of the GTP cavity colored according to electrostatic potential (from −15 kT/e, red, to +15 kT/e, blue) in open (**i**) and closed (**j**) conformations. **k**,**l**, The γ-phosphate coordinates a network of H-bond interactions with SW1 and SW2 residues, and intra-switch interactions in the open (**k**) and closed (**l**) states.

A small, hinged motion of SW1 residues Tyr^32^, Asp^33^ and Pro^34^ closes the active site forming the Michaelis complex (observed from 8 to 15 hours post-photolysis) (**Fig. 2b,c**). This movement appears to be anchored by Thr^35^ coordinating the Mg^2+^, Phe^28^ (which stacks against the purine base) and Asp^30^ forming an H-bond with ribose 2’-OH. The side chain of Tyr^32^ rotates 90° packing against Pro^34^ and reaches within H-bond distance to Gln^61^ (**Fig. 2d,e**); stacking of Tyr^32^ and Pro^34^ forms a tweezer that appears to constraint the motion of Gln^61^ (**Extended Data Fig. 2e**). In the closed state, the hydroxyl group of Tyr^32^ and the amide backbone of Thr^35^ make H-bonds with O3G and O2G, respectively; a net gain of two hydrogen bonds relative to the open state (**Fig. 2d,e**). At this location, the OH group of Tyr^32^ next to γ-phosphate’s O1G could render this atom more susceptible to nucleophilic attack by W_cat_. Of particular interest is the high mobility of Gln^61^ which can be seen adopting multiple conformers (**Fig. 2f**) (in both the open and closed states) to coordinate interactions with Tyr^32^, γ-phosphate’s O1G and W_cat_ (**Extended Data** Figs. 2f **I-IV and 2g V-VIII**). The position and coordination of the W_cat_ varies between the two states; in the open, it forms H-bonds with the carbonyl of Thr^35^, and the amide backbones of Gly^60^ and Gln^61^; whereas in the closed, H-bonds are mainly observed with the Thr^35^ carbonyl (**Fig. 2g,h**) and the side chain of Gln^61^, which reorients to coordinate W_cat_, a key step towards catalysis. Solvent extrusion in the closed state shrinks the GTP pocket (**Fig. 2i,j and Extended Data Fig. 2h,i**) increasing the positive charge around the γ-phosphate^33^. The Polder difference map shows a significant reduction in γ-phosphate occupancy suggesting that active site closure accelerates γ-phosphate hydrolysis^24^ (**Fig. 1d**). Analysis of SW1 and SW2 loops in the open and closed states reveals that the switches fold through a network of interactions with the γ-phosphate at its core (**Fig. 2k,l**); hence, γ-phosphate hydrolysis could trigger unfolding of both switches.

To quantify the hydrolysis process (14 hours post-photolysis) we calculated Polder difference maps (omitting GTP from the calculation) and measured the β-γ and β-α distances between their centers of density at 4.5 0 values (**Fig. 1c,d**). During the hydrolysis state, the electron density around O3B atom (compared to O3A) weakens (**Fig. 3a**); the β-γ phosphate distance lengthens, the β-γ dihedrals nearly eclipse, and the γ-phosphate dihedrals flatten, leading to γ-phosphate-O3B bond breaking (**Extended Data Fig. 3a-d**). Mg^2+^ coordination and H-bonds with the amide nitrogen atoms of Thr^35^ and Gly^60^ and the side chains of Gln^61^, Lys^16^ and Tyr^32^ temporarily stabilize the freed PO3^-^ species hindering its release (**Fig. 3b**); we identified electron density for two stabilized PO3^-^ enantiomers, including a Walden inverted species^34^ (**Fig. 3b-d**). As the reaction continues towards the product state, a polder difference map showed electron density for a PO4^3-^ ion (**Fig. 3e and Extended Data Fig. 3e**) which intriguingly locates at the W_cat_ site; however, its larger Van der Waals volume induces repositioning of SW1 and SW2 residues (**Extended Data Fig. 3f**). The product GDP state has no electron density for the γ-phosphate or SW1 and SW2 residues (33-37 and 61-68, respectively) suggesting that the energy released during GTP hydrolysis unfolds the signaling competent GTP-bound state (**Fig. 3f**) to unstructured switches in the GDP state incapable of interacting with effectors. The surface representation calculated for SW1 and SW2 regions during the hydrolysis and product states shows extensive remodeling, from a closed site (**Fig. 3g,h**) to formation of a short PO3^-^/PO4^-3^ exit tunnel (**Fig. 3i**), ending with unstructured switch loops (**Fig. 3j**). It is interesting to note that the Mg^2+^ ion shifts from its caged- and open-state position, where it is closer to β-phosphate’s O2B, towards the γ-phosphate’s O2G in the closed- and hydrolysis-state structures (**Fig. 3b-d**); returning to the β-phosphate O2B in the product state (**Fig. 3e,f,k and Extended Data Fig. 3m**). During release, the γ-phosphate dihedral angles follow a clockwise screw trajectory (**Fig. 3l,m**).

**Fig. 3:**
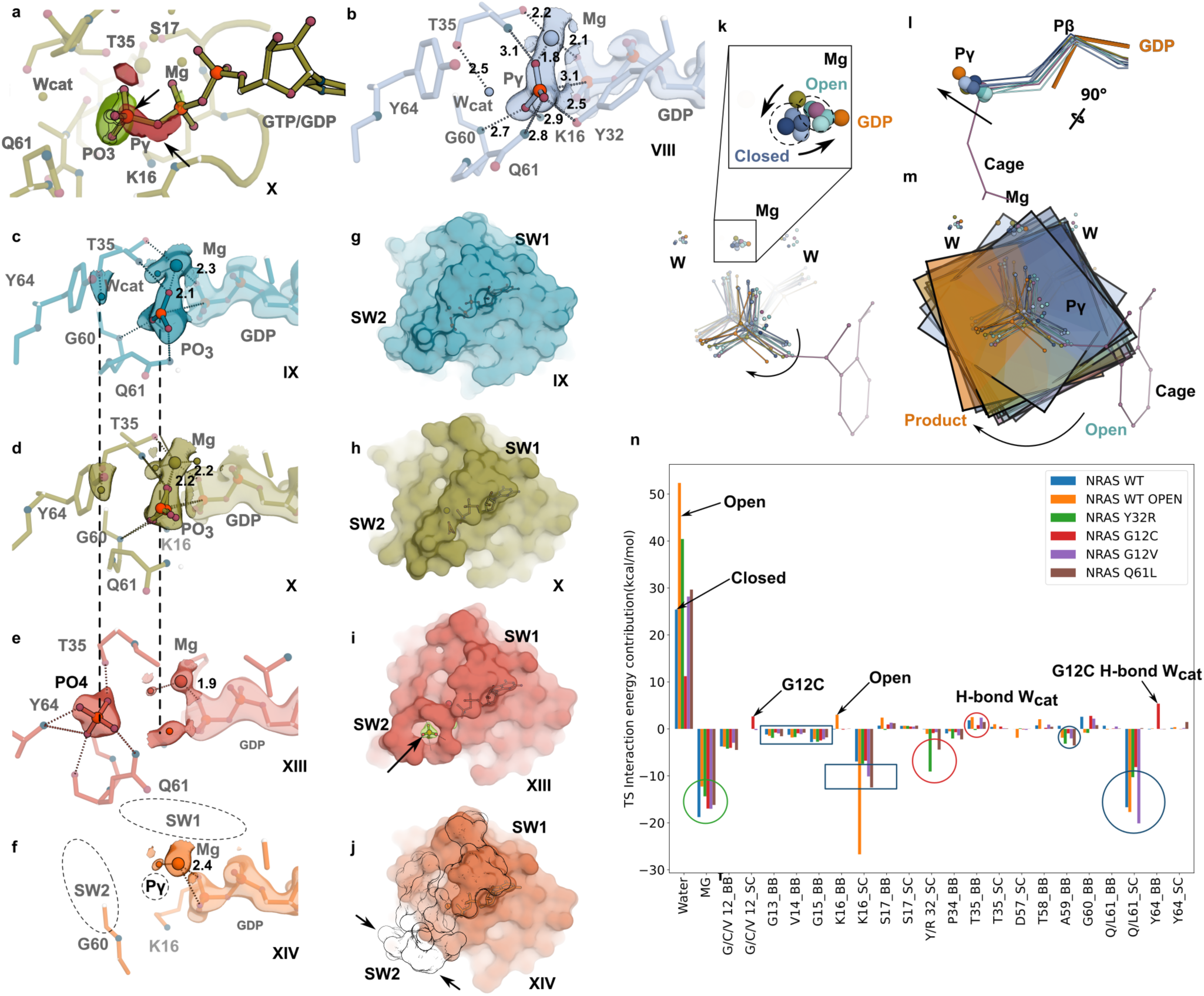
γ-phosphate Bond Breaking and PO3^-^/PO4^3-^ release and stabilization. **a**, *<u>Switch view.</u>* Difference maps contoured at +/- 30. Negative (red, arrow) and positive (green) density around O3B and γ-phosphate atoms illustrating β-γ bond breaking (negative density) and a position shift of the γ-phosphate (positive density) during the hydrolysis state. **b**-**f**, Ball and stick representation of the interactions stabilizing the freed PO3^-^/PO4^3-^ during the hydrolysis reaction. 2Fo-Fc electron density maps are shown as colored surfaces and are contoured above > 1.5 0 and B-sharpening of −15. Vertical dashed lines across (c-e) indicate the position of the γ-phosphate and Wcat. **g**-**j**, Surface representation of states IX, X, XIII and XIV illustrating formation of a short tunnel for PO3-/PO4^-3^ release (arrow and green density in **I**) and loss of the fully structured switch regions (black trace in **j**). **k**, State-colored spheres illustrating the changing positions of the Mg^2+^ ion (**k**) and the PO3-/PO4^3-^ (**l**) in the open, closed, hydrolysis and product states. **m**, γ-phosphate dihedral angles follow a clockwise screw trajectory during release. **n**, Chemical group contributions to the difference of interaction energies of W_cat_-GTP complex in the Transition State (TS) and initial state (Michalis complex). _BB and _SC denote backbone (amide) and side chain groups of protein residues, respectively. TS residue contribution. Green circle, Mg^2+^ ion; dark blue square, P-loop residues; red circle, SW1 residues; blue circle, SW2 residues.

## Computer Simulations

An EVB (Empirical Valence Bond) model^11,26,27^ was constructed to describe GTP hydrolysis via a solvent assisted mechanism based on DFT calculations of modeled hydrated phosphate compounds (see Methods). Analysis of the molecular dynamic (MD) trajectories of the activation free energy calculations allowed us to decipher the detailed molecular mechanism of the observed reaction rate changes. The computed hydrolysis rate for the open conformation shows a delayed reaction (**Extended Data Table 1**), highlighting the importance of the open-close conformational changes resulting in a smaller pocket around the GTP (**Fig. 2i,j**). Analysis of the molecular group contributions to the transition state (TS) (**Fig. 3n**) shows the large positive energy provided by water molecules, which results in screening of the electrostatic interactions around the GTP phosphate groups, and stabilization of W_cat_ in the initial reaction state. Hence, closing of the “reaction pocket” effectively expels water molecules around the GTP (**Extended Data Fig. 2h,i**), leaving the W_cat_ isolated and destabilized; thus, the transition from open to closed conformation effectively promotes the formation of the TS for the hydrolysis reaction. Another important aspect of the simulations shows that Mg^2+^ is better positioned to stabilize the TS in the closed conformation than in the open, as shown by its more negative contribution to the TS energy (**Fig. 3n**). The latter correlates with a shorter Mg^2+^-O2G distance in the closed state (**Fig. 3b-d**).

Evaluation of the energy contributions to the TS shows that the largest negative (stabilizing) energy comes from the Mg^2+^ ion, the Lys^16^ side chain, and the Gln^61^ side chain (**Fig. 3n**). Additional (stabilizing) energy contributions of individual amide backbone groups and amino acid side chains to the TS include: 1) amide backbone of P-loop residues 14-16, which form an open ring of positive charge; and 2) the backbone groups of SW1 Pro^34^ and SW2 Ala^59^ as these groups interact with the attacking water molecule in the TS geometry. Conversely, TS energy contributions from the backbone groups of SW1 Thr^35^, and SW2 Thr^58^ and Gly^60^ are positive (**Fig. 2g,h**) as these groups stabilize the attacking water in the initial (λ = 0) geometry of the reactive complex (**Extended Data Fig. 3n,o**).

## The Y32R mutant and the mechanism of hastened GTP hydrolysis

Structural overlay of N-RAS open and closed states with the K-RAS/RAS-GAP complex^14^ reveals that the arginine finger in RAS-GAP is positioned next to the γ-phosphate, but clashes with Tyr^32^ in the closed structures (**Extended Data Fig. 4a,b**). Analysis of the residues surrounding the γ-phosphate shows that the guanidinium group of the arginine finger (in the K-RAS/RAS-GAP structure) substitutes the hydroxyl group of Tyr^32^, hence increasing the positive electrostatic potential around the γ-phosphate (**Extended Data Fig. 4c,d**). To test whether such positive potential could have an impact on GTP hydrolysis, we constructed mutants where the negative charge of Tyr^32^ was replaced by positively charged residues (arginine, histidine or lysine) and thus mimic the function of the RAS-GAP arginine finger^14^. Only diffracting crystals of the N-RAS Y32R mutant bound to NPE-GTP were obtained. The caged structure shows the arginine residue with the aliphatic chain and the guanidinium group packing against the NPE moiety **(Extended Data Fig. 4e**). Overlay of both caged Y32R and WT structures shows they are nearly identical (r.m.s.d of 0.24 on Cα-carbons); given our expectation of a faster hydrolysis rate for this mutant, we pursued significantly shorter post-photolysis freezing times (**Fig. 4a**). The one-minute post-photolysis structure shows a closed active site, a poised arginine, with both guanidinium nitrogen atoms interacting with the γ- and α-phosphates, W_cat_ poised for nucleophilic attack and standard Mg^2+^ coordination (**Fig. 4b**). In the two-minute data set, Arg^32^ moves towards the γ-phosphate and Gln^61^ realigns to form an H-bond with a γ-phosphate oxygen, allowing arrival of the W_cat_ (**Fig. 4c,e**). Intriguingly, at this time point, the Mg^2+^ ion is found in two positions: the canonically coordinated site, and a second location (site B), slightly shifted away from the γ-phosphate (replacing one of the coordinating waters), and achieving an alternate coordination through Asp^57^ and the carbonyl oxygen of Val^58^ (**Fig. 4 c,e**). During this state, the Polder difference map shows a significant drop in γ-phosphate occupancy **(Extended Data Fig. 4f,g**). The five-minute time point shows the γ-phosphate out of the Mg^2+^ coordination sphere and rotated towards Arg^32^ disrupting H-bonding with Lys^16^ and Gly^60^. At this juncture, the Mg^2+^ is now fully positioned at site B (**Fig. 4d,f**), and the Polder map shows γ-phosphate occupancy below 40% **(Extended Data Fig. 4f,g**). By the ten-minute time point, the hydrolysis reaction is nearly complete (**Fig. 4 g,i,j)** with residual density for the γ-phosphate (occupancy is below 20%) and no density for Arg^32^ or Gln^61^; the Mg^2+^ ion is still found at the site B position. GTP hydrolysis is complete by the twenty-minute data point with unstructured SW1 and SW2 residues 32-37 and 60-68 (respectively) and the Mg^2+^ ion returning to its canonical position (**Fig. 4h,k**). Given the fast reaction rate, a single PO3^-^/PO4^3-^ intermediate with a slight β-γ phosphate bond-length increase was captured during the reaction (10 min) with the β-γ phosphate distance lengthening to 3.3 Å (**Extended Data Fig. 4h**). This appears to be yet another instance of a dynamical Mg^2+^ changing position during an enzymatic reaction^35,36^. Computer simulations illustrate that the TS stabilizing effect of the Y32R mutation is clearly seen in the large energy contribution of the Arg^32^ side chain (**Fig. 3n and Extended Data Fig. 4i,j**). GTP hydrolysis of the Y32R mutant parallels RAS-GAP reaction rates.

**Fig. 4:**
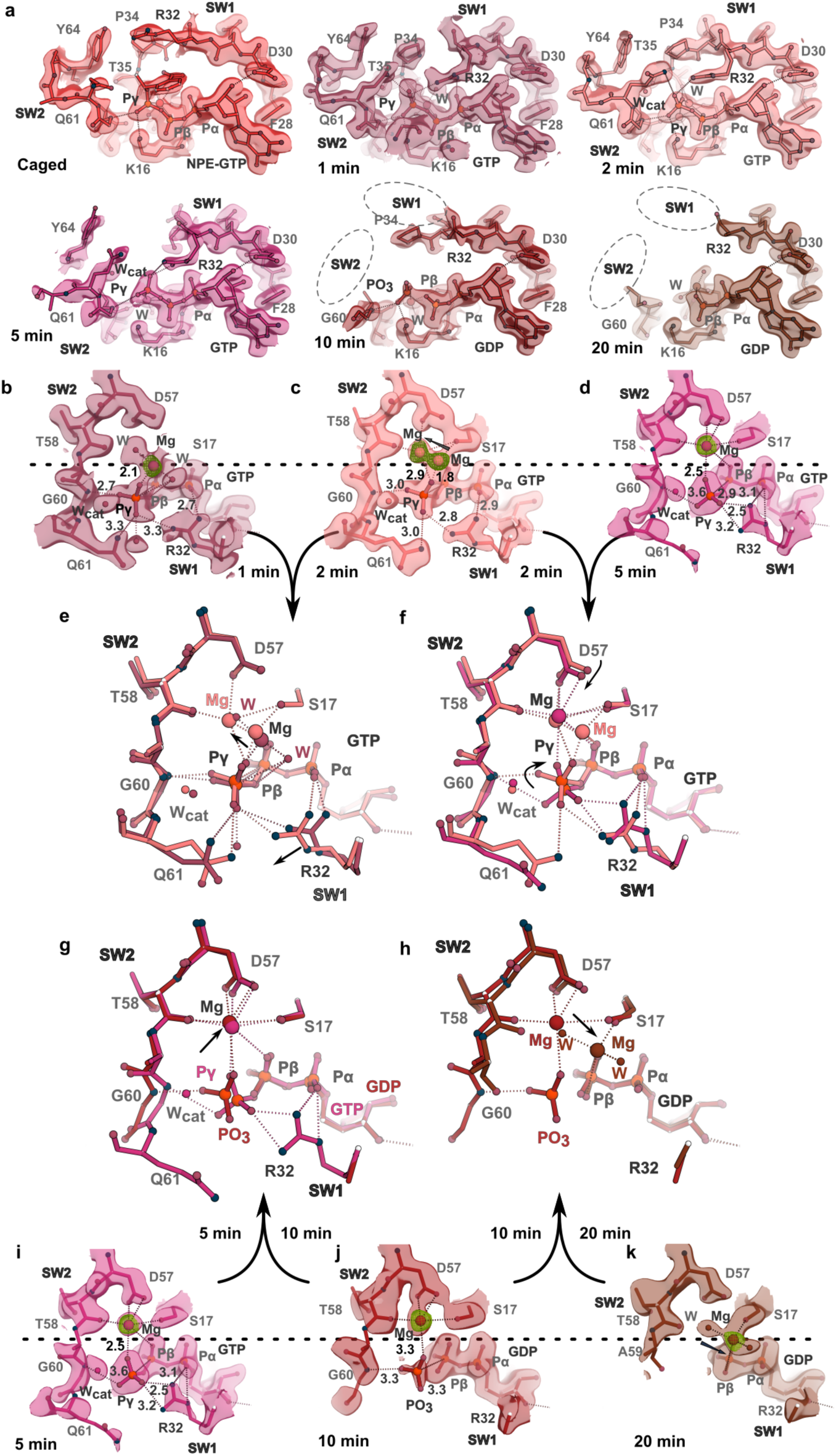
The structural basis of Y32R heightened GTP Hydrolysis. Palette of pink-colored surfaces illustrate 2Fo-Fc maps contoured at 1.5 0; green meshes are the Fo-Fc difference maps contoured at 3 0 (where Mg^2+^ was removed from the calculation). **a**, *<u>Switch</u> <u>view.</u>* Electron density maps and cartoon representation of the 6 reaction states (1min-20mins) for the Y32R mutant. *<u>Active site view</u>* illustrating Mg^2+^ coordination at the (**b**) 1-, (**c**) 2- and (**d**) 5-minute post-photolysis time points (notice that the γ-phosphate density at the 5-minute time point is diffuse, possibly indicating dynamical changes). The Mg^2+^ ion shifts (arrow in **c**) from the canonical position to a second one (site B) coordinated by the carbonyl of Thr^58^ and Asp^57^. **d**, The Mg^2+^ ion is now fully observed at site B; the dashed line across is drawn to illustrate the canonical position of the Mg^2+^ ion; undefined density of the 2Fo-Fc map around the γ-phosphate at the 5-minute time point could indicate mobility or more than one conformation (c.f. 1- and 2-minutes, where sharp density is found around the γ-phosphate). **e**, Cartoon and ball and stick representation of the overlay between the 1- and 2-min post-photolysis structures (red and light pink, respectively) showing Arg^32^ reaching towards the γ-phosphate. **f**, Overlay between the 2- and 5-minute (hot pink) structures; at the 5-minute time point Arg^32^ guanidinium group rotates and pulls the γ-phosphate away from Mg^2+^ coordination and H-bonds with Gly^60^. **g**, Overlay of the 5- and 10-minute time points showing Mg^2+^ ion at site B and SW1 and SW2 melting at the 10-minute time point. **h**, Overlay of the 10- and 20-minute time points showing the return of the Mg^2+^ ion to the canonical site and no density for the γ-phosphate or the switches. **i**-**k**, *<u>Active site view,</u>*electron density maps to illustrate Mg^2+^ coordination at the 5-, 10- and 20-minute post-photolysis time points. The Mg^2+^ ion stays at the alternate coordination position at the 5- and 10-minute time points, returning to its canonical position at the 20-minute time point.

## Functional Consequences of N-RAS Y32R/K/H mutations on cellular transformation

To understand the effect of the N-RAS Y32R/K/H mutants on the ability of N-RAS to mediate cellular transformation, we used a classic model of cellular RAS transformation in NIH 3T3 cells^37,38^. As the RAS Y32R mutant hydrolyzes GTP significantly faster, we hypothesized that it could prevent cellular transformation. We tested the effects of Y32R, and the two positively charged Y32K and Y32H mutations, on the ability of N-RAS oncogenic mutant G12C to transform NIH 3T3 cells. Typically, G12C oncogenic mutants trigger cellular transformation and senescence of NIH 3T3 cells upon DNA transfection or viral transduction, as well as increased net proliferation, by 96 hours. Interestingly, we found that evidence of transformation was still observed with the N-RAS Y32H mutant but transformation was not evident after infection with the N-RAS Y32K or Y32R mutants **(Extended Data Fig. 4k**). This suggests that the N-RAS Y32K or Y32R mutation are loss-of-function mutations. We next examined if a double N-RAS G12C/Y32R, G12C/Y32K or G12C/Y32H mutant, would prevent N-RAS G12C induced transformation. Of note, we found that transformation and induced senescence was still observed in the N-RAS G12C/Y32K and G12C/Y32H double mutants. Remarkably, the double mutant G12C/Y32R was unable to induce oncogenic transformation or senescence, which suggests that the Y32R mutation is sufficient to overcome oncogenic mutations in RAS **(Extended Data Fig. 4k**). We next attempted to quantitate the change in NIH 3T3 proliferation after transduction with these mutants **(Extended Data Fig. 4l**). As expected, the N-RAS G12C mutant showed a significant increase in proliferation compared to vector control. Intriguingly, none of the N-RAS Y32X mutants increased proliferation compared to the control vector, suggesting that these mutations are loss-of-function mutations as well. Remarkably, all three double G12C/Y32X mutants reversed G12C-induced proliferation **(Extended Data Fig. 4l**). In summary, these findings suggest that Y32R/K/H mutations are sufficient to overcome N-RAS-driven proliferation.

## Structural Basis of GTP hydrolysis of P-loop G12C, G12V and SW2 Q61L oncogenic mutants

Crystallization conditions for wild-type (WT) N-RAS with freely moving SW1 and SW2 residues were reproduced for oncogenic P-loop G12C and G12V, and SW2 Q61L mutants, enabling structural comparisons that were not crystal-contact biased. Overall, hydrolysis of the mutants proceeded appreciably slower, consistent with results of biochemical experiments^5,31,39,40^. Nineteen intermediate structures were captured for the G12C mutant (**Fig. 5a**). The time-resolved series (**Fig. 5a and Extended Data Fig. 5a-h**) showed a slower hydrolysis rate than WT (25-vs 15-hours respectively). The NPE-GTP structure shows an open active site with the thiol side chain of Cys^12^ pointing away from the γ-phosphate, and no density for SW1 and SW2 residues 35-37 and 61-67; hence, a water molecule replaces Mg^2+^ coordination by Thr^35^ (**Fig. 5b**). Comparison with the WT NPE-GTP structure shows Cys^12^ pointing away from the active site clashing with Gln^61^ and thus disrupting SW1-SW2 contacts (**Fig. 5e**). The open state is short-lived compared to WT; post-photolysis structures reveal that the active site closes within 2 hours (**Fig. 5a and Extended Data Fig. 5a,b**). Thr^35^ engages in Mg^2+^ coordination, and Cys^12^ rotates towards the γ-phosphate reaching within H-bond distance to O1G, Gln^61^ and Tyr^32^ **(Extended Data Fig. 5b).** Data points from 2 hours to 12 hours post-photolysis show well folded SW1 residues, however, SW2 residues Glu^62^ to Met^67^ remain unstructured (**Extended Data Fig. 5b,c**). Moreover, density for the W_cat_ is present but scarce, indicating low occupancy.

**Fig. 5:**
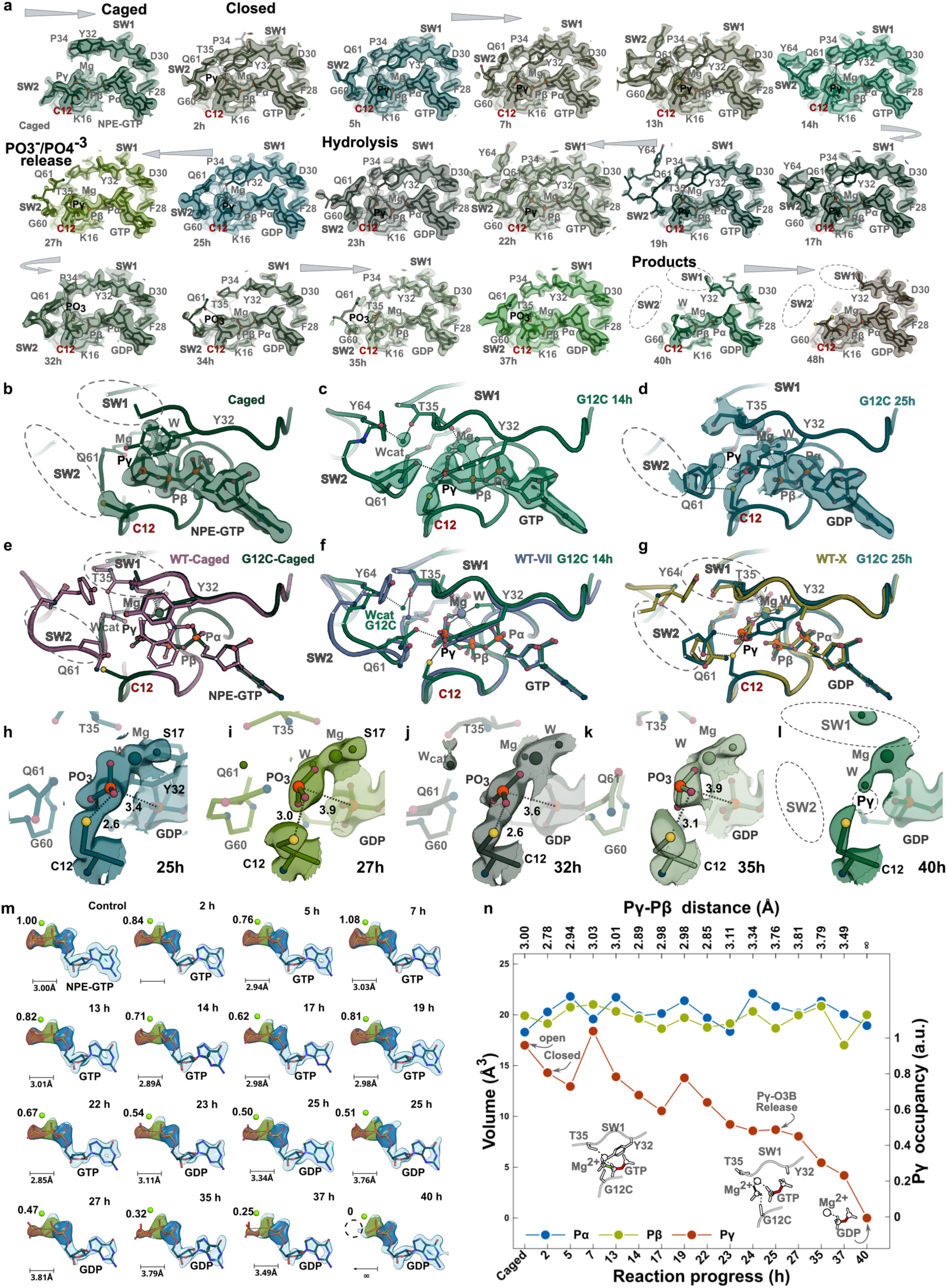
Structural Basis for γ-phosphate hydrolysis by the oncogenic G12C mutant. **a**, *<u>Switch view.</u>* The 2Fo-Fc maps are shown as surfaces colored using a green palette and contoured at 1.5 0 for eighteen GTP hydrolysis states. Transition from open to closed states occurs within the first 2-hours post-photolysis. Ball and stick representation (switch view) of selected time points: caged NPE-bound state (**b**), 14-hours post-photolysis (**c**), and 25-hours post-photolysis (**d**); electron density 2Fo-Fc maps are shown as colored surface contoured at 1.5 0. **e**-**g**, Overlay of WT and G12C structures for caged, 14- and 25-hours, respectively. **h**-**l**, Ball and stick representation of the interactions stabilizing the freed PO3^-^/PO4^3-^ during the hydrolysis reaction. The electron density map is contoured as colored surface at 1.5 0. Dashes depict H-bonds. **m**, Polder maps contoured at 4.5 0 omitting GTP from the calculation. Blue, green and red spheres correspond to α-β- and γ-phosphates, respectively. Bar illustrates the β-γ bond distance. **n**, Occupancy of the γ-phosphate as a function of the reaction process.

Structures from 14 to 19 hours show fully folded SW1 and SW2 residues, and Cys^12^ interacting with Tyr^32^, Gln^61^ and the γ-phosphate O1G (**Extended Data Fig. 5d,e**). It is interesting to note that coordination by the carbonyls of Thr^35^ and Tyr^64^ pulls W_cat_ away from the γ-phosphate (**Fig. 5c,f**), and thus the observed W_cat_-γ-phosphate distance is larger than WT (4.6A vs 3.5Å, respectively). The Pγ-Pβ distance increases around 22 hours post-photolysis; the 25-hour structure shows a free PO3^-^ forming multiple stabilizing H-bonds with Gln^61^, Gly^60^, Lys^16^, Thr^35^ and the thiol group of Cys^12^ (**Fig. 5d,g** and **Extended Data Fig. 5f**). As in the WT structure, two isomers of PO3^-^ were captured (**Fig. 5h-k**), and SW1 (32-37) and SW2 (61-68) residues unfold in the product state around 40 hours post-photolysis (**Fig. 5l**). The high mobility of Gln^61^ in WT structures is not observed in the G12C mutant since Cys^12^ and Tyr^32^ interactions constrain its motion (**Extended Data Fig. 5i-m**). Polder map analysis of time points spanning 25 to 40 hours shows a slow decay of γ-phosphate occupancy (**Fig. 5m,n**) possibly due to Cys^12^-OG1 interactions that slow down its release. In summary, our time-resolved data show a dynamic Cys^12^ adopting multiple conformers along the reaction trajectory and interacting with SW1 Tyr^32^, SW2 Gln^61^ and γ-phosphate O1G (**Extended Data Fig. 5n**).

MD simulations correlate with our structural observations and show that the G12C mutation destabilizes the TS of the reaction (**Fig. 3n**), as Cys^12^ side chain interacts with γ-phosphate’s O1G, stabilizing the initial reaction complex and hampering TS progression (where the O1G atom has a smaller negative charge). This is manifested in a positive energy contribution of Cys^12^ side chain to TS stabilization (**Fig. 3n**). One can also note a positive energy contribution of the backbone of Tyr^64^ to TS (**Fig. 3n**) due to stabilization of the attacking water in the initial reaction complex geometry (**Fig. 5c,f**).

The time-resolved series for the G12V mutant (**Fig. 6a and Extended Data Fig. 6a-i**) showed a significantly slower hydrolysis rate than WT or the G12C mutant. The NPE-GTP state bears some resemblance with the NPE-GTP state for G12C since no electron density for the NPE moiety, SW1 (35-37) and SW2 (65-68) residues was present (**Fig. 6b and Extended Data Fig. 6a,j**). Comparisons with the WT NPE-GTP structure shows Val^12^ clashing with the backbone of Gln^61^, positioning its side chain in the W_cat_ location and disrupting SW1-SW2 interactions (**Fig. 6b**). The open-state structures show Val^12^ turning towards the γ-phosphate, Gln^61^ rotating outwards allowing W_cat_ arrival, and Thr^35^ reaching the Mg^2+^ coordination sphere thus triggering SW1 folding (**Fig. 6c and Extended Data Fig. 6a,b**). The open state in G12V lasts significantly longer (16 hours) than in the native protein (∼8 hours) or G12C (∼2 hours), closing only partially, since electron density for Tyr^32^ is weak or not detected due to clashes with Val^12^ (**Extended Data Fig. 6c**). Bond lengthening, decreased γ-phosphate occupancy and unfolding of the switch loops starts around 48-hours (**Extended Data Fig. 6d-g**). By 72-hours the reactions is nearly complete with scattered density for the switch loops and minimal occupancy (**Fig. 6d,e and Extended Data Fig. 6h**). The product state is observed 96-hours post-photolysis with no discernable density for SW1 and SW2 loops (residues 34-37 and 59-69, respectively) (**Extended Data Fig. 6i**). As observed for Cys^12^ several rotamers of Val^12^ are seen in the time-resolved series (**Extended Data Fig. 6j,k).** Moreover, Gln^61^ shows lower mobility (when compared to WT N-RAS) owing to packing contacts with Val^12^ (**Fig. 6c,d and Extended Data Fig. 6k**)). While Val^12^ reaches within CHO bond distance to γ-phosphate O1G (**Extended Data Fig. 6j-48hrs**), it does not destabilize the TS as seen for Cys^12^ (**Fig. 3n**)

**Fig. 6:**
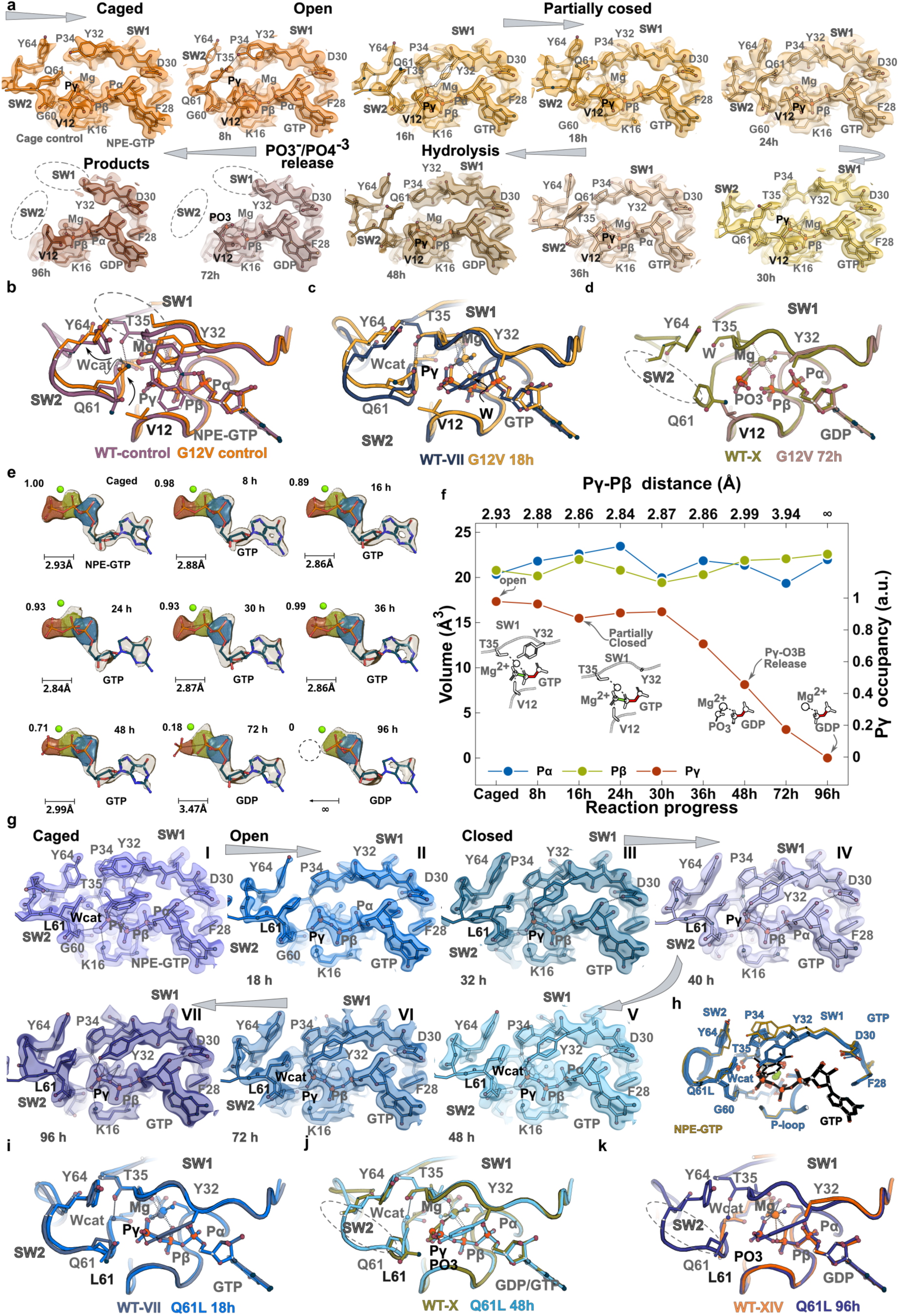
Structural Basis for γ-phosphate hydrolysis by the oncogenic G12V and Q61L mutants. **a**, *<u>Switch view.</u>* 2Fo-Fc maps shown as surfaces, colored using an orange-brown palette and contoured at 1.5 0 for ten GTP hydrolysis states; transition from open to a partially closed state occurs around 16 hours post-photolysis; however, no density for the side chain of Tyr^32^ is visible in the closed state. Cartoon and ball and stick representation of the overlay between the WT and G12V structures for the caged (**b**), open (**c**), partially closed (**d**) and PO3^-^/PO4^-3^release states. **e**, Polder maps contoured at 4.5 0 omitting GTP from the calculation. Blue, green, and red spheres correspond to α- β- and γ-phosphates, respectively. Bar illustrates the β-γ bond distance. **f**, Occupancy of the γ-phosphate as a function of the reaction process. **g**, 2Fo-Fc maps for the Q61L mutant shown as colored surfaces using a blue-violet palette and contoured at 1.5 0 for the seven GTP hydrolysis states; transition from open to closed states occurs between the 18-32 hours post-photolysis. **h**, Overlay of Q61L structures I-VII illustrating the “frozen” nature of the intermediate states; the NPE-caged structure is shown in gold. Cartoon and ball and stick representation of the overlay between the WT and Q61L structures for caged (**i**), 48-hours (**j**), and 96 hours (**k**) illustrating the presence of folded switches for the Q61L mutant in all time points.

The Q61L structures show well-defined caged, open and closed states (**Fig. 6g,h**); however, as expected^5,39^, GTP hydrolysis was slowed down significantly in the crystals. The NPE-GTP structure in Q61L is remarkably similar to the WT structure with a r.m.s.d on Cα_s_ of 0.38 (**Extended Data Fig. 6l**). Transition from the open to the closed state occurs between 18 and 32-hours post-photolysis since WT Gln^61^-Tyr^32^ interactions promoting active site closure are not present, hence Tyr^32^ is dynamic, impacting its ability to form a H-bond with γ-phosphate’s O1G. During the closed state, a stalled Michaelis complex is formed but does not lead to productive hydrolysis. Analysis of the Polder maps shows full occupancy of the γ-phosphate for the first 72 hours, reaching 40% occupancy at 96 hours post-photolysis (**Extended Data Fig. 6n,o**) and the β-γ phosphate bond length remains constant throughout the reaction increasing slightly at 96 hours post-photolysis. Q61L structures are characterized by small motion of Leu^61^, decreased mobility of SW1 and SW2 residues (**Extended Data Fig. 6p**), and a nearly fixed W_cat_ (**Fig. 6i,j**). Structural overlay with WT N-RAS shows minimal conformational changes with nearly “frozen” structures in eight states (**Fig. 6i-k**). Moreover, in contrast with the P-loop Cys^12^ and Val^12^ mutants, no rotamers of Leu^61^ are observed in the time-resolved series (**Extended Data Fig. 6m).**

To rationalize the observed changes of GTP hydrolysis rates in N-RAS mutants, we performed detailed molecular simulations of their reaction mechanisms. Here we rely on the findings that GTP hydrolysis in RAS proteins proceeds via a “solvent-assisted” mechanism^11^ with the rate limiting step being the formation of the dissociative TS with the water molecule attacking the metaphosphate-like intermediate (**Extended Data Fig. 3m**) (see methods) without the need of a general base. The results for the activation free energy calculations are shown in **Extended Data Table 1**. Computed changes of GTP hydrolysis rates in WT N-RAS compared to the mutants qualitatively reproduce the effects observed in the structures (**Extended Data Table 1, 3^rd^ column**); including two-orders of magnitude increase in GTP hydrolysis rate for N-RAS Y32R; and a strong decrease in the hydrolysis rate for Q61L mutant, underscoring its role in organizing the electrostatic environment of the reaction center including Tyr^32^, the γ-phosphate and W_cat_ interactions.

## Discussion

GTP and ATP hydrolysis are highly regulated events, critical to signaling pathways that impact a wide variety of cellular functions, including proliferation, cell trafficking, nuclear transport and cell motility. In particular, the RAS family of small GTPases are critical regulatory proteins controlling cellular proliferation and their oncogenic mutants are present in a large number of malignancies^1^. To gain new insights into the reaction intermediates during GTP hydrolysis, we postulated that N-RAS bound to the photoactivated NPE-GTP could allow for precise temporal control of the reactants; and that crystallization conditions with unconstrained SW1 and SW2 mobility could enable observation of GTP hydrolysis *in crystallo*. Generating crystals that diffracted to high-resolution (1.6-1.9 Å) enabled observation of 57 reaction intermediates and hence, the conformational changes during GTP hydrolysis for native N-RAS, the fast-hydrolyzing Y32R, and the oncogenic P-loop G12C, G12V, and SW2 Q61L mutants; the hydrolysis mechanisms differed for each oncogenic mutant and proceeded at slower rates than native N-RAS.

The NPE-moiety in GTP generated a new crystalline state, effectively hindering the hydrolysis reaction for both N-RAS and K-RAS. However, crystal contacts trapping SW1 Tyr^32^ in K-RAS crystals lead to a non-catalytic configuration (**Extended Data Fig. 1d**). Conversely, following cage release, N-RAS residues in the open state adopt a pre-catalytic configuration where SW1 and SW2 loop convergence form a H-bond network with the γ-phosphate at its core; the folded switches create a small pocket that traps W_cat_ (achieving coordination with the carbonyl of Thr^35^); and Gln^61^ positions to interact with the γ-phosphate and the W_cat_. The pre-catalytic conformation leads to the catalytic state during active site closure. The long-lasting open state is intriguing since the closed structure (Michaelis complex) should be reached relatively fast following cage release at 16 °C. This is true for Y32R where the active site is closed at 1-min post-photolysis (illustrating that the conformation induced by the NPE-cage is not restrictive); however, it is not the case for WT, G12C, G12V and Q61L mutant structures, where it persists for 8 to 11-, 2-, 18- and 32-hours (respectively), in a “locked” open state. Moreover, our structural data (**Fig. 1d**) and simulation experiments show that GTPase activity is inhibited in this conformation (**Extended Data Table 1**). Desolvation of the active site has been shown to favor enzyme catalysis^29,41^; hence, conversion between the pre-catalytic to the catalytic conformation in N-RAS might rest on the ability to desolvate the GTP binding pocket during active site closure. Our simulations point to the large stabilizing energy contributed by the solvent (**Fig. 3n**); it is possible that overcoming the large positive energy provided by water molecules might be energetically costly leading to a prolonged open state.

The biological relevance of a sustained open state may rest on the observation that N-RAS interactions with the RAS-GAP could be more favorable with open state structures, since the arginine finger (Arg^789^) clashes with Tyr^32^ in the closed but not the open conformation (**Extended Data Fig. 4a,b**). Conversely, overlay with structures of the B-RAF^42^, PI3K^43^ and Sin1^44^ effectors, illustrate that these molecules can bind to both the open- and closed-conformations (**Extended Data Fig. 7a-f**). Since WT open state structures have 8-fold lower hydrolysis rates (**Extended Data Table 1**), RAS-GAP binding and hydrolyzing open state structures could limit the number of longer-lived activated RAS molecules; yet, closed state structures would still inactivate through intrinsic GTP hydrolysis. On the other hand, it is possible that RAS-GAP Arg^789^ could displace Tyr^32^ triggering GTP hydrolysis; however, insertion of the arginine finger seems to be critical for complex formation as we were unable to assemble a N-RAS-NF1 complex in the presence of NPE-caged GTP, since Arg^789^ clashes with the NPE group.

Detailed molecular simulations of the TS formation provided additional details of the importance of closing the GTP pocket for the reaction by promoting stability of the TS, and the effects of amino acid mutations on the reaction rates (**Extended Data Table 1 and Fig. 3n**). Conformational changes that promote stability include: 1) γ-phosphate O1G-Tyr^32^ H-bond formation; 2) active site volume contraction and extrusion of water molecules to promote isolation and destabilization of W_cat_ for γ-phosphate nucleophilic attack; 3) Gln^61^ coordination of W_cat_; and 4) Mg^2+^ ion drifting towards the γ-phosphate (**Extended Data Fig. 3m**), weakening the Pγ-O3B bond^33^. These conformational changes correlate with a loss in γ-phosphate occupancy and lengthening of the β-γ-phosphate distance (**Fig. 1d and Extended Data Fig. 3q**-v).

We capture electron density resembling PO3^-^, a mixture of PO3^-^/PO4^3-^ with Walden inverted^34^ PO3^-^ species and a released PO4^3-^ trapped by switch residues for the hydrolysis state structures (**Fig. 3b-e**). The occupancy of an atom in a crystal is given by the fraction of asymmetric units that contain it. In the caged state, since the NPE-cage group inhibits hydrolysis, the occupancy for α-β- and γ-phosphates should be close to one. Following NPE photolysis, the fraction of asymmetric units containing the γ-phosphate changes as the reaction evolves from the early open states (close to 1) to the product GDP state (effectively 0) (**Fig. 1c,d**). Since GTP hydrolysis by N-RAS is an intrinsically slow process, three populations of N-RAS molecules could coexist (in the crystal) during the reaction: 1) molecules that have not undergone hydrolysis, (i.e., have an intact GTP) that will be found predominantly in the open state; 2) N-RAS molecules with a hydrolyzed γ-phosphate observed in the closed, hydrolysis and release states; and 3) N-RAS molecules with a GDP substrate, seen primarily at the end of the reaction. Thus, the observed electron density for the PO3^-^/PO4^3-^ species is the superposition of these three populations, and is mainly determined by the occupancy of the γ-phosphate, since the B-factor values for the β- and γ-phosphates remain close to each other while the difference in occupancies increases significantly (**Extended Data Fig. 3p**). Contributing elements to the stability of the released PO3^-^/PO4^3-^ species include multiple H-bonds with neighboring residues and Mg^2+^ coordination (**Fig. 3b-d,n)**. Computational studies demonstrated the stability of a PO3^-^ intermediate and suggested its universal presence in all nucleotide hydrolysis events^45^.

In contrast to most K-RAS, N-RAS and H-RAS structures reported to date, the presented GDP (product) states in WT, G12C, G12V, and Y32R mutants have unfolded SW1 and SW2 loops; two functional implications could be drawn from these findings. First, since SW1 and SW2 loops create the effector binding surface, the unfolded switches of the GDP state cannot sustain effector interactions **(Extended Data Fig. 7a-f**) ^46^. Second, the unfolded switches could facilitate binding to the guanine nucleotide exchange factors (GEFs) (**Extended Data Fig. 7g,h**).

The effect of the Y32R mutant during GTP hydrolysis, while expected, is still noteworthy since positioning a guanidinium group next to the phosphate tail mimicking the arginine finger increases the reaction rate by 135-fold. The open state could not be detected in Y32R mutant possibly as a result of the strong Arg^32^ interactions with the α- and γ-phosphates^47^. However, consecutive time points show Arg^32^ moving towards the γ-phosphate and positioning its guanidium group for catalysis. Moreover, the TS snapshot (λ = 12) of the Y32R mutant in our simulations shows that the side chain of Arg^32^ reorients to coordinate the O3B and O2A atoms of the leaving GDP group (**Extended Data Fig. 4i,j**), contributing to stabilization of additional negative charge accumulated (on the α- and β-phosphate groups) during γ-phosphate-O3B bond breakage. The high resolution of the structures allowed observation of the dynamics of the Mg^2+^ ion during the hydrolysis reaction where it is found in two positions. First, the canonical site contributing to hydrolysis (**Fig. 4b**); the second (site B), with γ- and β-phosphates out of range of its coordination sphere would favor PO3^-^/PO4^3-^ release (**Fig. 4i,j**). The latter could be a result of the large electrostatic pull^47^ from Arg^32^ guanidium group on the freed phosphate. Hence, the combined effect of charge stabilization and its release from the Mg^2+^ coordination sphere in the Y32R mutant could be the two main contributing factors to its fast hydrolytic rate.

Our time-resolved structural and modeling data show that reaction mechanisms and rates for the native and each of the mutants differ significantly. Comparisons of the WT and mutant structures reveal differences in the GTP hydrolysis mechanism that correlate with the computed hydrolysis rates in our simulations (**Extended Data Table 1 and Extended Data Fig. 7i,j**). The time evolution of the G12C structures and the computed reaction times show that GTP hydrolysis proceeds at a slower rate than WT. Three possible mechanisms could account for this observation. First, although the closed state is significantly shortened by Cys^12^-Tyr^32^ interactions (two hours post-photolysis), SW2 residues (62-67) are disordered and the solvent around W_cat_ stabilizes the initial reaction state, hence slowing down hydrolysis. Moreover, the mobility of Gln^61^ is restricted by its interactions with Cys^12^ and Tyr^32^ (**Extended Data Fig. 5i-l**). Second, although folding of SW2 residues around the 14-hour time point generates the Michaelis complex; however, differences in the arrangement of SW2 residues with respect to WT places Tyr^64^ carbonyl within H-bond distance of W_cat_ pulling it 1.0 Å farther from the γ-phosphate, hampering catalysis (**Figs. 5c,f** and **3n)**. Third, Cys^12^ interactions with the freed PO3^-^/PO4^3-^ (25-hour time point) delay its release as illustrated by its slow occupancy decay (**Fig. 5n**) and its positive contribution to the TS (**Fig. 3n**). Our structural data and molecular simulations show that a mobile Gln^61^ sidechain adopts different conformations in the reaction stabilizing the attacking water in native N-RAS; conversely, the G12C mutation interferes with Gln^61^ rearrangement and stabilizes the initial state of the reaction, slowing hydrolysis. The presence of bulky side chain at the P-loop tip was proposed to inhibit RAS-GAP/NF1 inactivation due to potential clashes with the arginine finger^48^. Overlay of the G12C (14-hours)-RAS-GAP structures^14^ show steric clashes of the arginine finger with both Cys^12^ and Tyr^32^ (**Extended Data Fig. 5o,p**) precluding interactions with GAP proteins.

The G12V P-loop mutant shows significantly more profound effects for the hydrolysis reaction. Active site closure was observed at longer times (approximately 18 hours post-photolysis). The observed delay could be related to absence of Tyr^32^-O1G or Tyr^32^-Gln^61^ contacts due to steric clashes with Val^12^ hampering active site closure and extrusion of water molecules from the active site pocket. Our molecular simulations (**Extended Data Table 1**) show that hydrolysis rates for the open state of the WT protein are comparable to the G12V mutant. GTP hydrolysis starts at the 48-hour time point and finalizes around the 96-hour time point (**Extended Data Fig. 6g-i**).

The Gln^61^ mutant showed the slowest rate of hydrolysis in the structural as well as the simulations, highlighting its importance in providing the electrostatic environment towards hydrolysis. Our results illustrate the mechanism of Gln^61^ in GTP hydrolysis and its role in organizing the electrostatic environment of the reaction center, including Tyr^32^ phosphate and W_cat_ interactions. The mutant shows a nearly identical open state with the WT structure (r.m.s.d 0.38) and was characterized by a decreased mobility of SW1 and SW2 residues. The transition to the closed conformation occurred between the 18- and 32-hour time points; this late transition is possibly a consequence of the lack of Leu^61^ and Tyr^32^ interactions. The results of our simulations can also explain the important role played by Gln^61^ in GTP hydrolysis since its side chain in the transition state trajectory (λ=12) (**Extended Data Fig. 3m,n**) changes orientation to coordinate and stabilize the attacking water molecule, hence explaining the reduction in the hydrolysis rate in the N-RAS Q61L mutant whose side chain cannot interact with the attacking water.

P-loop NTP-ases comprising the GTPase family (including small-GTPases and the Gα-subunit of the heterotrimeric-GTPases) and ATPases (including molecular motors such as myosin and kinesin) feature a conserved structural motif (Walker-A motif) around the NTP substrate, comprising P-, SW1- and SW2-loops^49^. We have shown that the variability in hydrolysis rates – from 5 minutes to more than 96 hours– arises from residue composition of the Walker-A motif. Thus, while most RAS Walker residue types are similar in the Gα family, the conserved SW1 Arg^178^ (equivalent to NRAS Tyr^32^) enhances the reaction rate (as observed for the Ras-GAP and the Y32R mutant); however, it is slower than expected^50^ (2 min^-1^) since its position is modulated by P-loop residue Glu^43^ which pulls it away from the γ-phosphate (**Extended Data Fig. 8a,b**). Comparably, the basic features of the GTPase hydrolysis reaction appear to be preserved for the ATPases kinesin and myosin: 1) both ATPases feature an asparagine residue in SW1 (equivalent to NRAS Tyr^32^ or Arg^32^) which H-bonds with α- and γ-phosphate oxygens and has been proposed to play a critical role in hydrolysis^51,52,53^ (**Extended Data Fig. 8c,d**); 2) the transition state structures of both myosin and kinesin^54,55^ show stabilizing interactions through Mg^2+^ coordination and H-bonds with P-loop, SW1 and SW2 residues, which closely resemble WT intermediate state IX (**Extended Data Fig. 8e-h**); 3) Our studies suggest a critical role of the γ-phosphate to assemble and sustain an effector-binding conformation with folded switches (**Fig. 2k,l**). Myosin gate residues Arg^243^ (SW1) and Glu^466^ (SW2) form a salt bridge creating a barrier to PO4^3-^ release; as is the case for our WT VII-XI structures, myosin pre-power stroke structures show a trapped PO3^-^/PO4^-3^ species in the active site stabilized by neighboring P-SW1- and SW2-loop residues and coordinated by Mg^2+56^. MD simulations show a potential release pathway with PO3^-^/PO4^-3^ binding sites in the myosin tunnel^56^. However, no structure to date has captured the PO4^-3^ ion the tunnel, since release structures show an open gate with ADP. Analogous to our state XIII structure, where the released PO4^-3^ ion occupies the position of the W_cat_, myosin residues stabilizing the W_cat_ create a charged pocket that could seize the released PO4^-3^ bounded by gate residues (**Extended Data Fig. 8i**).

Kinesin and myosin exploit the energy released during γ-phosphate hydrolysis to perform mechanical work; however, the significance of energy release for GTPases has not been elucidated. Our observation that the switches are unstructured in the GDP state might indicate that the energy released during GTP hydrolysis disrupts the structured (effector-binding) switches, swiftly ending the signaling event.

Traditional structure-based drug screens for RAS GTPases are restrictive since non-hydrolysable analogs are employed, and molecules in the crystal reach steady state well before analysis at the synchrotron. Structure-based drug discovery has mostly relied on “static snapshots’’ but not experimentally resolved intermediates whose targeting can also impact biological function. Search for cryptic binding surfaces during GTP hydrolysis in G12C, G12V, and Q61L mutants reveals the presence of several (unique) state-dependent binding pockets, hitherto inaccessible for drug discovery (**Extended Data** Figs. 5q-s and 6p**,q**). Employing RAS proteins with “live” GTP molecules could allow enhanced sampling of the conformational space for small fragment binding in oncogenic mutants.

## Methods and Materials

### Cloning, Expression and Purification

N-Ras in pET28 vector was a gift from Cheryl Arrowsmith (Addgene plasmid # 25256; http://n2t.net/addgene:25256; RRID: Addgene_25256). The mutants G12C, G12V, Q61L and Y32R were created by site-directed mutagenesis.

Cells expressing a 6-His-tagged N-RAS were resuspended in buffer A (100 mM NaCl, 25 mM Hepes pH 7.5 and 5 mM β-mercaptoethanol) and sonicated. Cell lysates were centrifuged and the supernatant was loaded onto a 5 mL Nickel-NTA column. Two washes of 20 column volumes each with buffer A containing 25 mM and 40 mM Imidazole, respectively, were followed by an elution step with buffer A containing 150 mM imidazole. The eluted sample was concentrated and dialyzed overnight in buffer A to remove imidazole. While maintaining a dark environment, 100 μM protein sample was mixed with 2.4 U of alkaline phosphatase, 10 mM EDTA and 5 mM NPE-caged GTP for 3 hours at 25 °C to hydrolyze the GDP molecule and load the caged GTP onto N-RAS. Following the incubation period, 10 mM MgCl_2_ was added to the sample and a size exclusion chromatography step with Buffer A plus 5 mM MgCl_2_ on a Superdex75 column was followed by sample concentration to a final concentration of 40 mg/ml.

### Protein Crystallization

Initial crystals of N-RAS were obtained in 20% PEG8K and 100 mM sodium acetate pH 5.6-6.0. Crystals were subjected to several rounds of micro-seeding to improve quality. To photolyze the NPE cage, crystals were exposed with light at 365 nm for 30 seconds, followed by freeze-quenching in liquid nitrogen at different intervals post-photolysis. Cryoprotection of crystals was accomplished by mixing a solution containing mother liquor plus 25% glycerol and 10% 2-methyl-2,4-pentanediol (MPD) as cryoprotectants.

### Cage Photolysis

A LED light source emitting at 365-nm with 1.2 W radiant flux was used for photolysis. The divergent light beam was shaped by a series of lenses such that the spot cross-section at the sample was (8 mm). The light was pulsed and its duration controlled by software.

### Structure Refinement and Analysis

The structures were solved by molecular replacement in MOLREP ^57^using as initial model PDB:ID 3CON. Model building and refinement were performed using Coot ^58^ and Phenix^59^. Figures with rendered using the PyMOL Molecular Graphics System, Version 3.0 Schrödinger, LLC. The structures have been deposited in the Protein Data Bank (see Data Availability)

### Calculation of electron density volumes around the α-, β-, and γ-phosphates

To quantify the local electron density specifically around the α-, β-, and γ-phosphates of the GTP ligand, we generated Polder difference maps (omitting GTP from the calculation) using the corresponding N-RAS structure for each system. From this initial map, the volume of electron density was measured independently for each of the three phosphate atoms. Using Coot v. 0.9.8.96^60^, a 2 Å radius mask was generated centered on each phosphate on the polder map. This masked map was then exported to UCSF ChimeraX v. 1.8^61,62^ for direct volume calculation. At an isovalue of 0.5 e-/Å³ (approximately 4 sigma), the enclosed volume for each phosphate was determined. Furthermore, the occupancy (probability of finding an atom at a particular location) of the γ-phosphate was assessed across the various systems by normalizing its calculated volume against the volume in the control system, providing a quantitative measure of its relative density.

**Diffraction Data Collection and Analysis** (see Supplementary Materials)

### Cell lines and reagents

The NIH 3T3 cell line was procured from American Type Culture Collection (ATCC) and grown in media recommended by ATCC.

### Lentivirus production, transduction, and preparation of stable cell lines

The lentiviral particles were created using a four-plasmid system and infected according to the TRC Library Production and Performance Protocols, RNAi Consortium, Broad Institute^63^, and as described previously^64,65^. Briefly, 2.5 x 106 HEK 293T cells were seeded in T-25 flasks, following a 24 h post incubation period, cells were transduced with respective plasmid DNA with the help of polyethylenimine (PEI) to generate lentiviruses. These vectors were used as template for a two-step PCR to add the remaining C-terminal amino acid sequence of N-RAS to these constructs and subsequently Gateway cloning was utilized to clone these inserts into the lentiviral vector (W118). All constructs were sequence verified and expression confirmed by western blot analysis.

### Cell transformation and viability assays

For cellular transformation assays, NIH 3T3 cells were seeded at a cell density of 3.0 × 10^5^ cells/well in a six well plate and incubated in 5% CO2 at 37^○^ C in a humidified atmosphere for 24 h. Images were obtained after 72 hours at 20X magnification. For all cell viability experiments, NIH 3T3 cell lines were seeded in 96 well plates at a cell density of 5.0 × 10^2^ cells/well and incubated in 5% CO2 at 37^○^ C in a humidified atmosphere for 24h. Cells were subsequently infected with the indicated vector or N-RAS mutant lentiviral constructs. Following 24 hours of infection, lentivirus was replaced with normal media or media with TKIs for 48 or 72 h. At 96 hours after infection, cell viability was assessed using CellTiter96 Aqueous One Solution Cell Proliferation Assay Kit (Promega) or CellTiter-Glo (Promega) according to the manufacturer’s protocol. Data was analyzed using the Graphpad Prism Version 8.4.3 software, as previously described^64,65^.

### Computational methods

An Empirical Valence Bond model (EVB)^11,26,27^ was constructed to describe GTP hydrolysis via a solvent assisted mechanism based on DFT calculations of modeled hydrated phosphate compounds. The EVB model was used to define the geometry of the transition state (TS) of the reaction, which is characterized by: 1) the 2.26 Å distance between GTP γ-phosphate and oxygen of the attacking water molecule P_γ_-W_cat_,; 2) the 2.56 Å length of the P_γ_-O3B bond that will be broken during the reaction. We computed activation free energies of the GTP hydrolysis performing alchemical transformations of H_2_O + Mg^2+^-GTP Michaelis complex to the TS geometry as defined above. The initial geometries of the reaction complexes for each N-RAS mutant were taken from the experimental structures obtained before visible GTP hydrolysis was observed for the WT N-RAS open (8 hours) and closed conformations (15hours). Corresponding time points were taken for Y32R, G12C, G12V and Q61L mutants at 2 minutes and 15, 30 and 72 hours, respectively.

To perform molecular dynamics simulations along the reaction coordinate Mg^2+^-GTP protein complexes were solvated in a water box of 56 x 56 x 75 Å, with Na^+^ cations added to ensure charge neutrality. The initial geometries of the simulated systems were subjected to 1000 steps of the steepest decent energy minimization followed by 30 ns relaxation MD simulations with restrained heavy atoms. Alchemical transformations of H_2_O + Mg^2+^ + GTP Michaelis complex-to TS-model used 13 intermediate state replicas with protein CA atoms restrained to experimental positions. Calculations for each replica were run for 200 ns, with the last 100 ns of each replica’s MD simulations used for free energy calculations, employing Thermodynamic Integration (TI) and Bennet Acceptance Ratio (BAR) methods.

## Data Availability

The atomic coordinates and structure factors have been deposited into the Protein Data Bank and will be available upon publication under accession numbers: 9PNZ (*NRAS WT-caged*), 9PO0 (NRAS WT-I), 9PO1 (NRAS WT-II), 9PO2 (NRAS WT-III), 9PO3 (NRAS WT-IV), 9PO4 (NRAS WT-V), 9PO5 (NRAS WT-VI), 9PO7 (NRAS WT-VII), 9PO8 (NRAS WT-VIII), 9PO9 (NRAS WT-IX), 9POA (NRAS WT-X), 9POB (NRAS WT-XI), 9POC (NRAS WT-XII), 9POD (NRAS WT-XIII), 9POE (NRAS WT-XIV), 9POF (*NRAS Q61L-caged*), 9POG (NRAS Q61L-18h), 9POH (NRAS Q61L-32h), 9POI (NRAS Q61L-40h), 9POJ (NRAS Q61L-48h), 9POK (NRAS Q61L-72h), 9POL (NRAS Q61L-96h), 9POM (*NRAS G12C-caged*), 9PON (NRAS G12C-2h), 9POO (NRAS G12C-5h), 9POP (NRAS G12C-7h), 9POQ (NRAS G12C-13h), 9POR (NRAS G12C-14h), 9POS (NRAS G12C-15h), 9POT (NRAS G12C-17h), 9POU (NRAS G12C-19h), 9POV (NRAS G12C-22h), 9POW (NRAS G12C-23h), 9POX (NRAS G12C-25h-1), 9POY (NRAS G12C-25h-2), 9POZ (NRAS G12C-27h), 9PP0 (NRAS G12C-32h), 9PP1 (NRAS G12C-35h), 9PP2 (NRAS G12C-37h), 9PP3 (NRAS G12C-40h), 9PP4 (NRAS G12C-48h), 9PP5 (*NRAS G12V-caged*), 9PP6 (NRAS G12V-8h), 9PP7 (NRAS G12V-16h), 9PP8 (NRAS G12V-18h), 9PP9 (NRAS G12V-24h), 9PPA (NRAS G12V-30h), 9PPB (NRAS G12V-36h), 9PPC (NRAS G12V-48h), 9PPD (NRAS G12V-72h), 9PPE (NRAS G12V-96h), 9PPF (*NRAS Y32R-caged*), 9PPG (NRAS Y32R-1min), 9PPH (NRAS Y32R-2min), 9PPI (NRAS Y32R-5min), 9PPJ (NRAS Y32R-10min), and 9PPK (NRAS Y32R-20min).

## Acknowledgements.

The contents of this publication are solely the responsibility of the authors and do not necessarily represent the official views of NIGMS or NIH. G.C. acknowledges support from R01GM112686 and R01AI175067. D.S. acknowledges support from the European Union’s Horizon 2020 research and innovation programme under the Marie Skłodowska Curie grant agreement No 884104 (PSI-FELLOW-III-3i). Use of the Stanford Synchrotron Radiation Lightsource, SLAC National Accelerator Laboratory, is supported by the U.S. Department of Energy, Office of Science, Office of Basic Energy Sciences under Contract No. DE-AC02-76SF00515. The SSRL Structural Molecular Biology Program is supported by the DOE Office of Biological and Environmental Research, and by the National Institutes of Health, National Institute of General Medical Sciences (P30GM133894). GM/CA@APS has been funded by the National Cancer Institute (ACB-12002) and the National Institute of General Medical Sciences (AGM-12006, P30GM138396). This research used resources of the Advanced Photon Source, a U.S. Department of Energy (DOE) Office of Science User Facility operated for the DOE Office of Science by Argonne National Laboratory under Contract No. DE-AC02-06CH11357. We thank Drs. Daniel Altschuller and Fatema Bhinderwala for their critical reading of the manuscript. G.C. would like to thank Dr. Michael Becker GM/CA@APS for assistance with data collection. We thank Prof. Cheryl Arrowsmith for the N-Ras plasmid

## Author Contributions

G.L., P.Z.P., X.Z., B.C., and D.C. performed expression and protein purification. X.Z. cloned NRAS WT and the G12C, G12V, Q61Land Y32R,H,K mutants. G.L., DC, BC and P.Z.P. performed crystallization experiments. G.L., S.R., M.E.S, D.P.S. and G.C. collected diffraction data. U.S. and G.C. performed the X-ray data processing and analysis. I.K. and M.G.K. performed molecular dynamic simulations. S.A.H., R.K.G. and T.F.B. performed cell transformation and viability assays with N-RAS, G12C and Y32R,H,K mutants. V.N. provided the optical system for cage photolysis. S.V. and A.D. contributed to the analysis. G.C conceived the project. G.C. and A.C design X-ray experiments and together with T.F.B., M.G.K., coordinated and supervised the work. G.C wrote the manuscript with input from all co-authors.

## Extended Data

**Extended Data Fig. 1:**
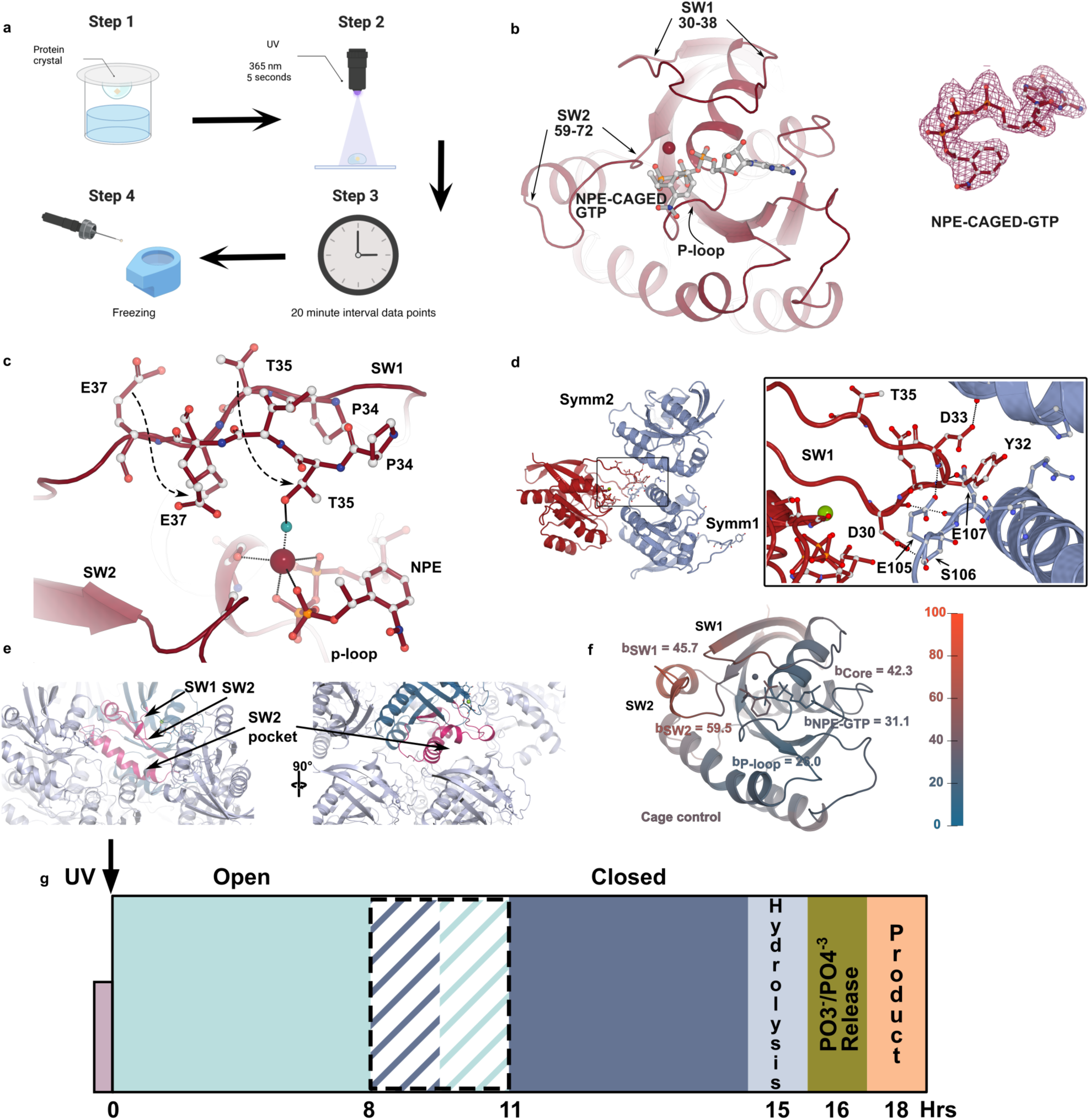
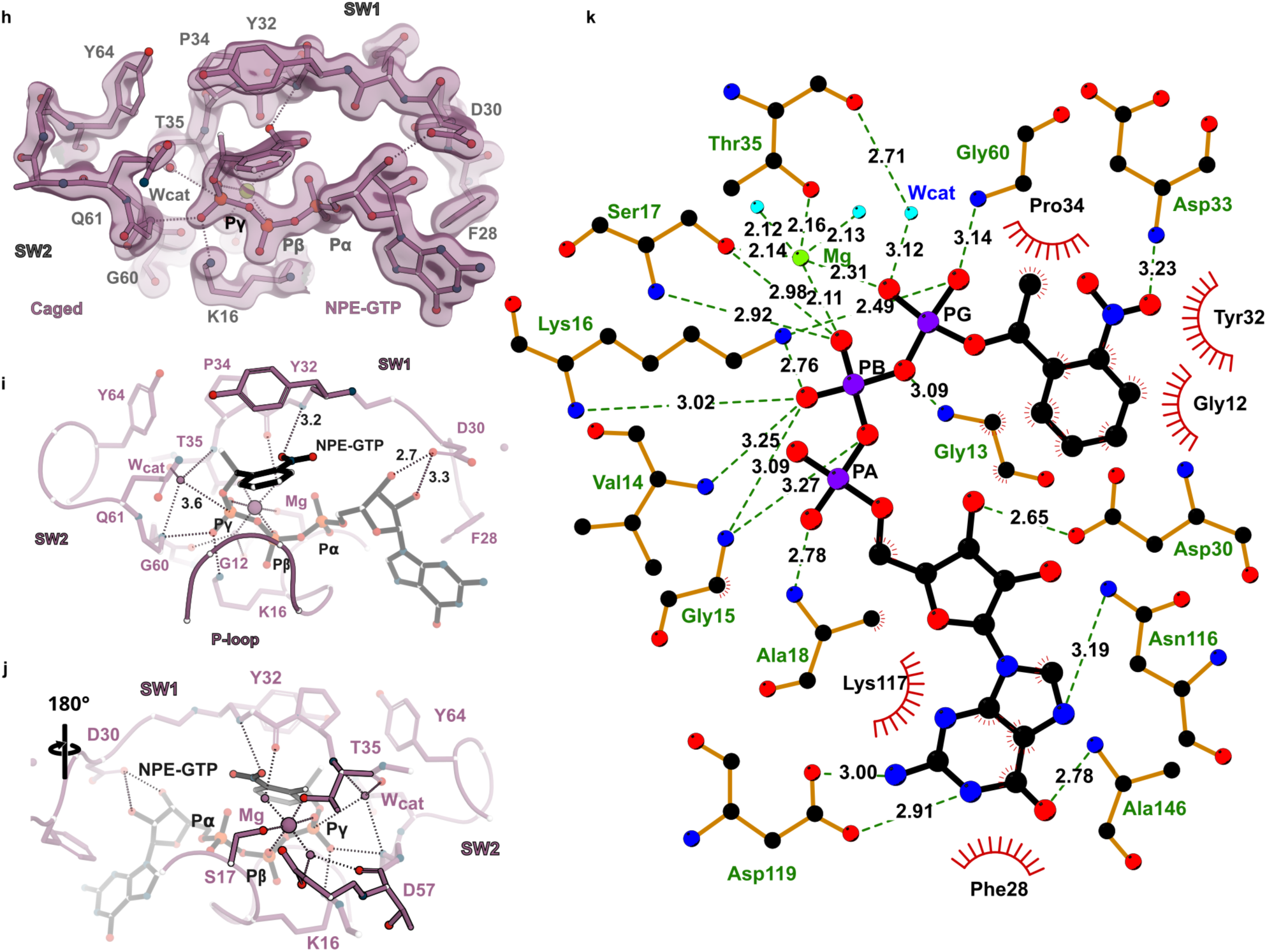
K-RAS and N-RAS NPE-caged X-ray studies. **a**, Experimental setup for UV-photolysis of “caged” NPE-bound K-RAS and N-RAS. **b**, Left panel, cartoon and ball and stick representation of K-RAS NPE-GTP-bound. Right panel, red mesh, Fo-Fc map corresponding to NPE-GTP contoured at 3.0 0. **c**, Two conformations of SW1 residues observed in the crystals. **d**, Crystal contacts of K-RAS SW1 with symmetry related molecules (colored in light blue). Inset illustrates trapping of SW1 Tyr^32^ inside a small pocket formed by Glu^105^, Ser^106^ and Glu^107^ and an adjacent helix. **e**, Cartoon representation of N-RAS (colored in blue) and its symmetry related molecules (colored in light blue); arrows indicate the position of SW1, SW2 and the SW2 pocket (colored pink), where effectors and drugs bind. **f**, Calculation of B-factors for N-RAS shows higher values for the switch regions as compared to the core, showing enhanced motility. **g**, Schematic of reaction progress during GTP hydrolysis; dashed lines indicate overlap between the states. **h**, Final refined 2Fo-Fc electron density map contoured at 1.5 0, (switch view); density for the NPE-cage can be easily identified. **i**,**j**, Two views of the cartoon and ball stick representation illustrating cage-NPE interactions in the pocket and canonical Mg^2+^ ion coordination by Ser^17^, Thr^35^, the β- and γ-phosphate oxygens and two waters molecules. Dashes represent H-bonds, and numbers distances in Å. **k**, Ligplot^66,67^ representation of the NPE-GTP interactions in N-RAS active site pocket.

**Extended Data Fig. 2:**
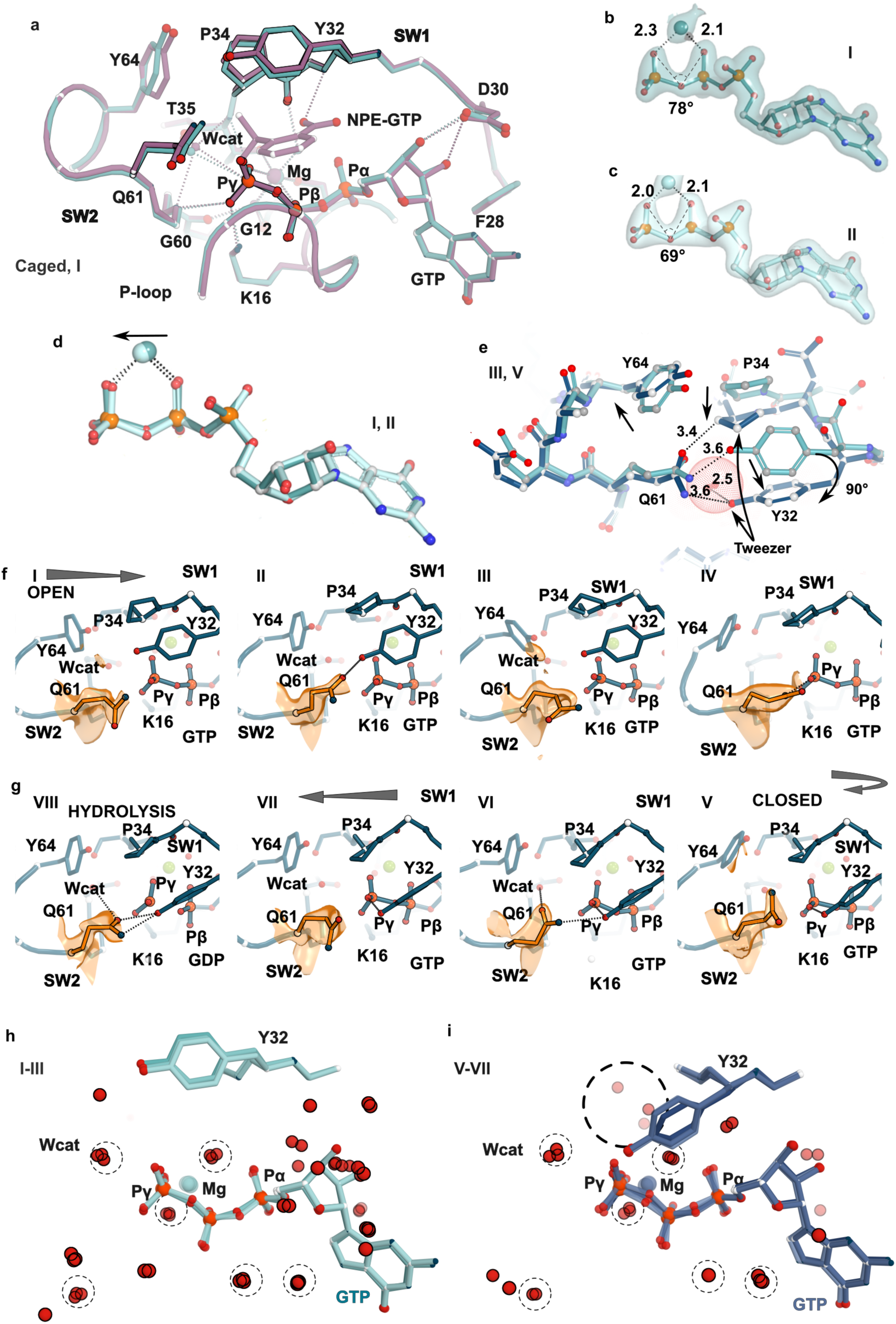
Transition from open-to closed-states. **a**, *<u>Switch view</u>* Overlay of the NPE-GTP (violet) and post-photolysis structure I (cyan) showing high similarity. Dashes represent H-bonds with distances below 3.5 Å. **b**, Ball-and-stick representation of GTP for open state I showing that the Mg^2+^ ion is closer to the β-phosphate, while a later time point (**c**, state II) shows that it has shifted towards the γ-phosphate. The Polder difference maps in **b** and **c** are shown as surface and contoured at 3 0 with –10 B-factor sharpening. **d**, Overlay of the ball and stick representation of GTP between open states I and II. The arrow illustrates the motion of the Mg^2+^ ion towards the γ-phosphate. Dashes illustrate distances below 3.5Å. **e**, Overlay of residues Tyr^32^, Gln^61^, Pro^34^ and Tyr^64^ between the open (cyan, state III) and closed (blue, state V) structures. Note the packing contacts between Pro^34^ and Tyr^64^ for both states, and the 90° side chain rotation of Tyr^32^ (in the closed state, arrow) to engage the γ-phosphate (shown as a red sphere). **f**, Mobility of Gln^61^ during the open (I-IV) and (**g**) closed states (V-VIII). Orange surface corresponds to the 2Fo-Fc contoured at 1 0 with a B-factor sharpening of −10. Arrows show the direction of the reaction. **h**,**i**, Representative solvent (water molecules) around the γ-phosphate in the (**h**) open (states I-III) and (**i**) closed states V-VII, respectively. Waters with similar positions in both states are surrounded by small dashed circles and include mainly W_cat_ and waters coordinating the Mg^2+^ ion. Water molecules cluster inside a small pocket formed by the side chains of SW1 residues in the closed state (large dashed circle).

**Extended Data Fig. 3:**
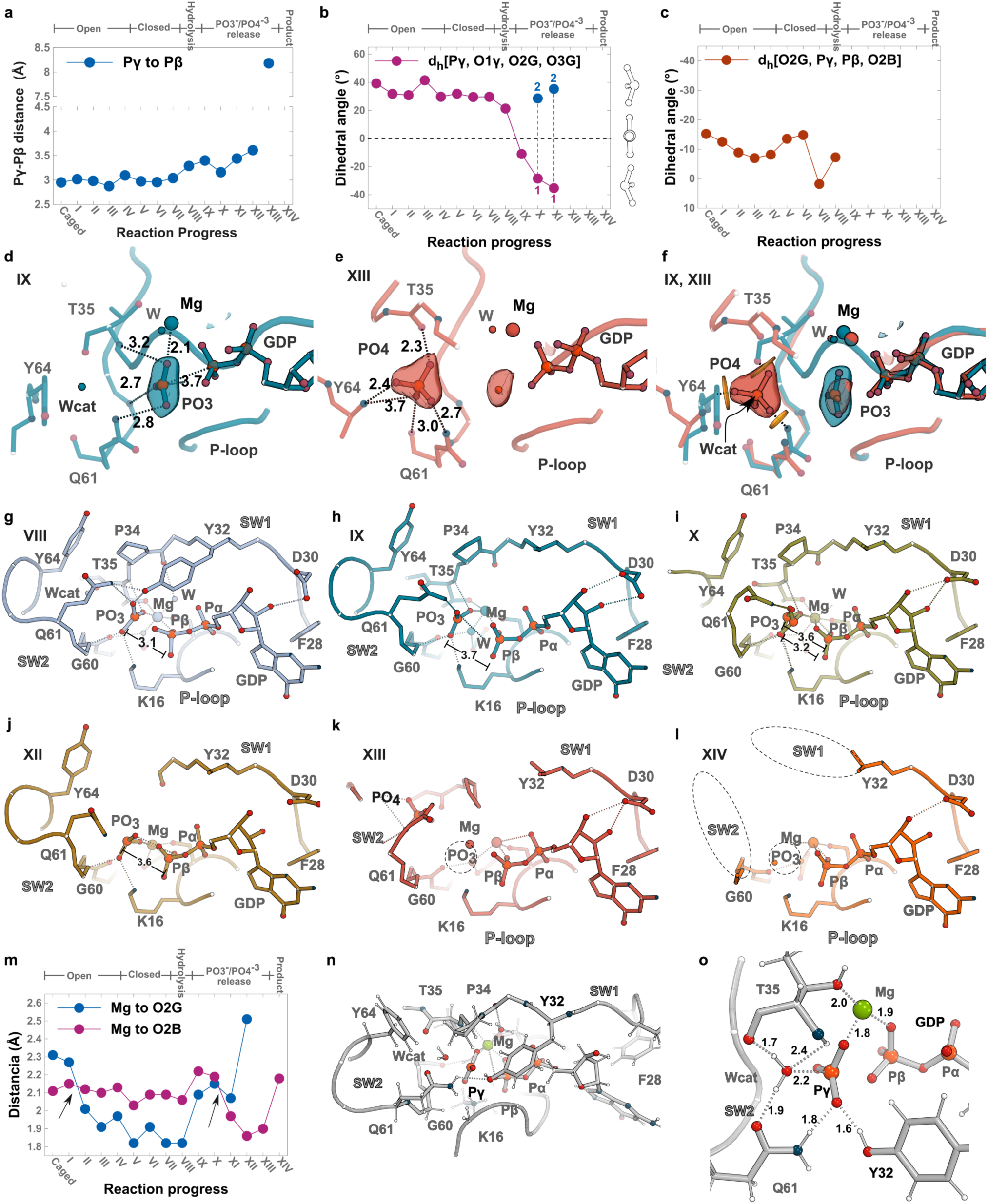
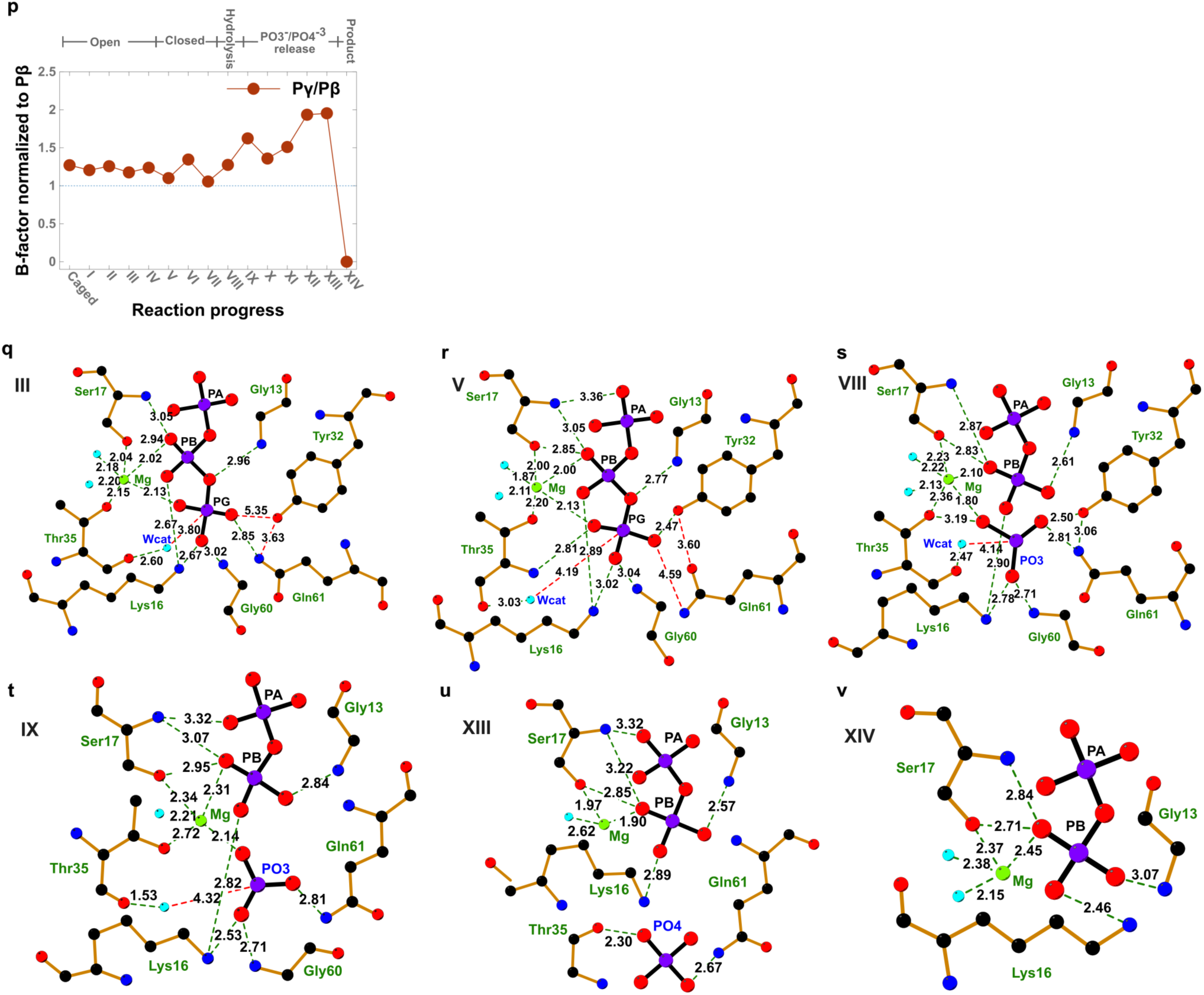
Bond lengthening and PO3-/PO4-3release. **a**, Reaction progress vs β-γ bond length. **b**, Reaction progress vs γ-phosphate dihedral angle (states X and XI have two enantiomers labeled 1 and 2). **c**, Reaction progress vs β-γ dihedral angles. **d**, *<u>Switch view</u>* illustrating a stabilized nearly flat metaphosphate (state IX, see also Extended Data Fig. 3b); the Polder difference map, contoured at 30, is shown as blue surface. **e**, Stabilization of the released PO_4_^3-^ ion through H-bonds with the carbonyls of Thr^35^ and Gln^61^, the amide backbone of Tyr^64^, and the side chain of Gln^61^; the Polder difference map, contoured at 30, is shown as red surface. **f**, Overlay of states IX (blue) and XIII (red) illustrating how clashes of state IX structure (yellow disks) displace SW1 and SW2 residues to allow PO_4_^3-^ binding. **g**-**l**, Cartoon and ball-and-stick representation (*<u>Switch view</u>*) of (selected) states illustrating the structural changes during γ-phosphate hydrolysis, comprising β-γ phosphate lengthening in Å (bars) and melting of the SW1 and SW2 loops; the GDP substrate state (**l**) shows fully unfolded switches. **m,** Mg^2+^-O2B (pink trace) vs Mg^2+^-O2G (blue trace) distance during the hydrolysis reaction. Arrows show crossing points. **n,o**, Cartoon and ball-and-stick representation of the transition state complex snapshot (colored silver) from Free Energy calculations (λ=12 trajectory). Gln^61^ side chain reorients to coordinate the attacking water. **o.** Reaction progress vs Pγ/Pβ B-factor. **p**-**v**, Ligplot representation illustrating contacts between α-β-γ phosphates and neighboring residues for selected open, closed, hydrolysis, PO_3_^-^/PO_4_^3-^ release and product states.

**Extended Data Fig. 4:**
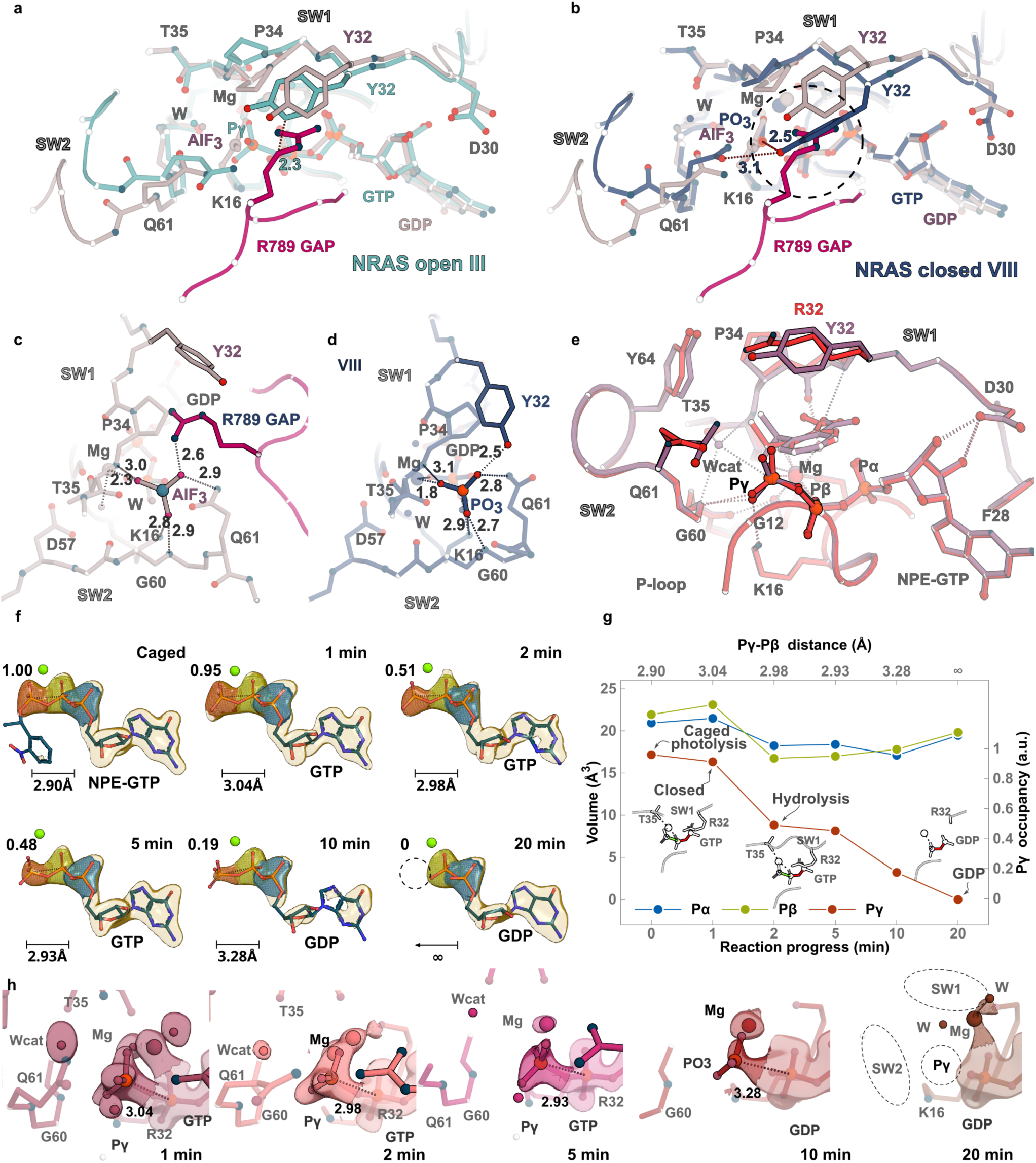
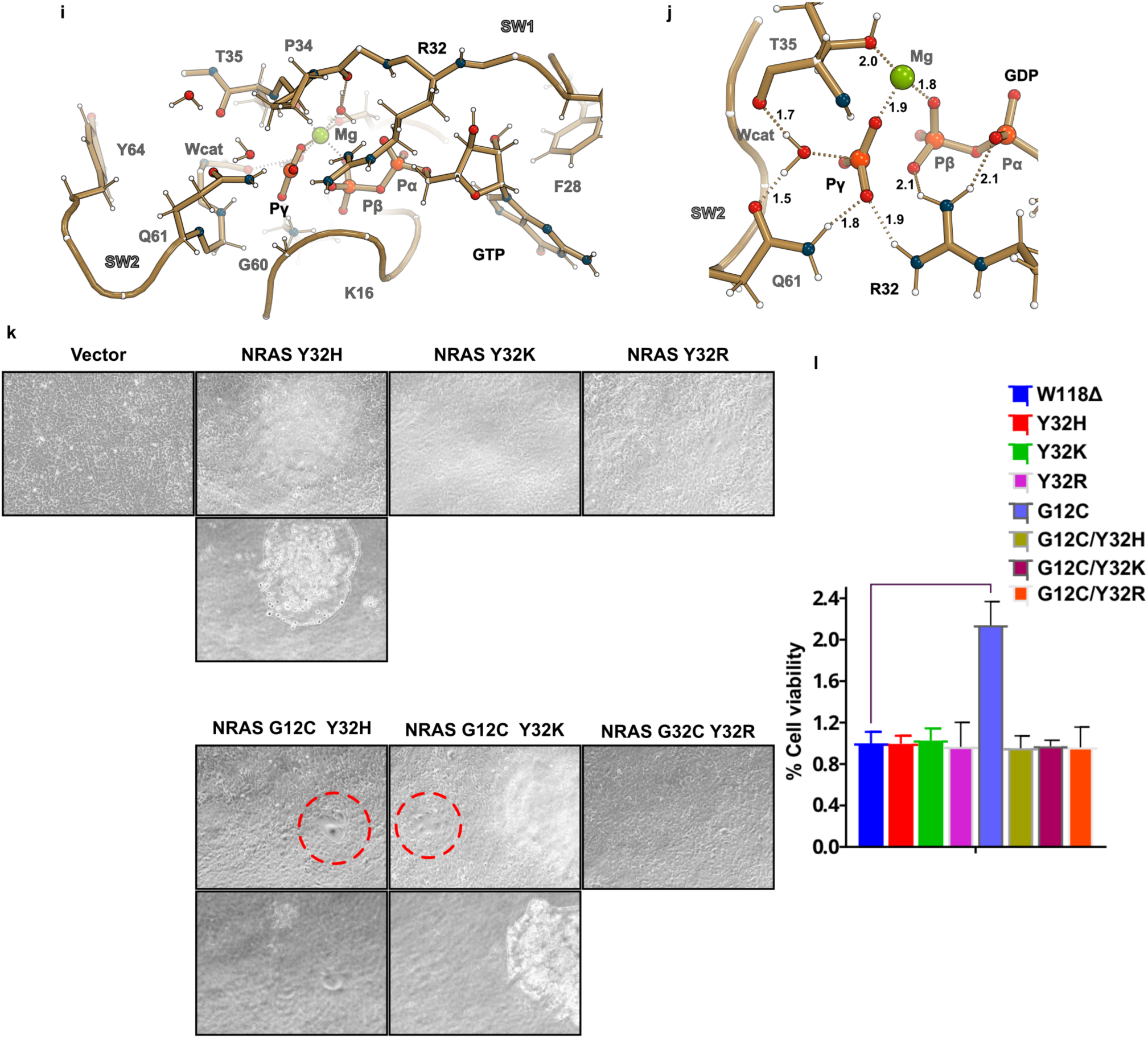
Y32R is a fast hydrolytic mutant. **a**,**b**, *<u>Switch view.</u>* Overlay of the ball and stick representations for K-RAS (light brown) RAS-GAP (dark pink) structure (PDB:ID 1WQ1) and the N-RAS (a) open, state III (cyan) and (**b**) closed, state VIII (dark blue) structures illustrating potential clashes between RAS-GAP Arg^789^ (arginine finger) and Tyr^32^ in the closed state (dashed circle). **c**,**d**, *<u>γ-phosphate view</u>* H-bonds between the γ-phosphate oxygen atoms and neighboring SW1, SW2 and P-loop residues for (**c**) K-RAS (light brown)/RAS-GAP (dark pink) complex (PDB:ID 1WQ1) and (**d**) N-RAS state VIII. Dashed lines indicate distance in Å. **e**, Switch view of the overlay between the N-RAS WT (light purple) and Y32R (red) NPE-GTP bound (caged structures). **f**, Polder maps contoured at 4 0 omitting GTP from the calculation. Blue, green and red spheres correspond to α-β- and γ-phosphates, respectively. Bar illustrates the β-γ bond distance. **g**, Occupancy of the γ-phosphate as a function of the reaction process. **h**, 2Fo-Fc electron density maps contoured at 2 0 around the GTP for 1-, 2-, 5-, 10- and 20-minute time points. **i**,**j** Transition state complex snapshot from free energy calculations (λ=12 trajectory) of NRAS Y32R mutant. Arg^32^ side chain reorients to coordinate O3B atom on the leaving phosphate group. **k**,**l**, NRAS mutants with increased hydrolysis prevent NRAS G12C mutant driven transformation and proliferation. **k**, Representative images (20X magnification) 72 hours after infection with the indicated vector control or NRAS mutant lentiviral constructs. Evidence of colony formation is reflected by multiple images taken at distinct focus points and red circles denote area containing senescent cells. **l**, Representative cell viability assay at 96 hours after infection of NIH 3T3 cells in quadruplicate demonstrated a significant increase in proliferation (p value 0.0006) in the NRAS G12C mutant compared to vector control (W118Δ) which is abrogated by the Y32X mutation. Of note, no other comparisons were statistically different.

**Extended Data Fig. 5:**
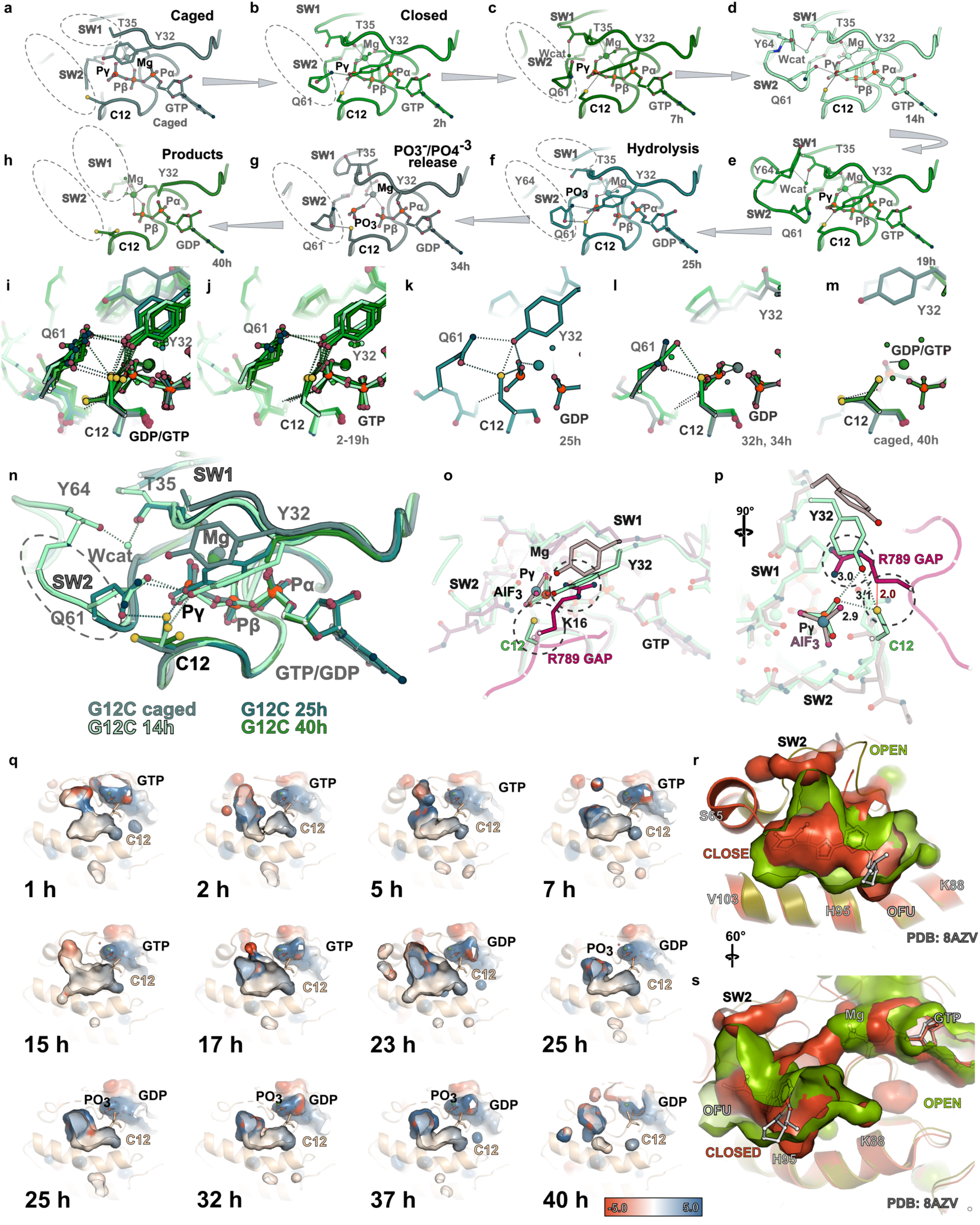
Prolonged PO3-/PO4-3release of the G12C mutant. **a**-**h**, Cartoon and ball and stick representation (*<u>switch view</u>*) of eight representative states (from 2-hours to 40-hours) during G12C GTP hydrolysis. Dashed ovals represent unfolded switch regions; note switch loops unfold/fold at different time points during the GTP hydrolysis reaction. **i**-**m**, Ball and stick representation (*<u>switch view</u>*) of Tyr^32^, Cys^12^, Gln^61^ and γ-phosphate’s O1G interactions during the indicated time points. **n**, Cartoon and ball and stick representation of Cys^12^ rotamers during the hydrolysis reaction. **o**,**p**, Two views (*<u>switch</u>* and *<u>γ-phosphate views</u>*, respectively) of the overlay between K-RAS/RAS-GAP (PDB:ID 1WQ1, hot pink) and N-RAS G12C structure (light green, 14-hours) illustrating potential clashes between Cys^12^ and Arg^789^, which could limit enhanced RAS-GAP GTP hydrolysis. Dashed circles indicate residue clashes. **q**, Surface electrostatic representation (colored according to electrostatic potential (from −5 kT/e, red, to +5 kT/e, blue) of the intermediate state SW2 cryptic pockets illustrating morphology changes at different times during GTP hydrolysis; **r**, Comparison between the SW2 cryptic pockets of the K-RAS *<u>GDP</u>* (PDB:ID 8AZV)^68^ (orange) and the N-RAS G12C *<u>GTP</u>* structure(s) (23 hours, green) illustrating the enhanced volume available for inhibitor binding. A modelled K-RAS inhibitor OFU ((4S)-2-azanyl-4-methyl-4-[3-[4-[(1S)-1-[(2S)-1-methylpyrrolidin-1-ium-2-yl]ethoxy]pyrimidin-2-yl]-1,2,4-oxadiazol-5-yl]-6,7-dihydro-5H-1-benzothiophene-3-carbonitrile) bound to the K-RAS GDP structure is colored in silver for illustration purposes and is shown as a ball and stick^68^.

**Extended Data Fig. 6:**
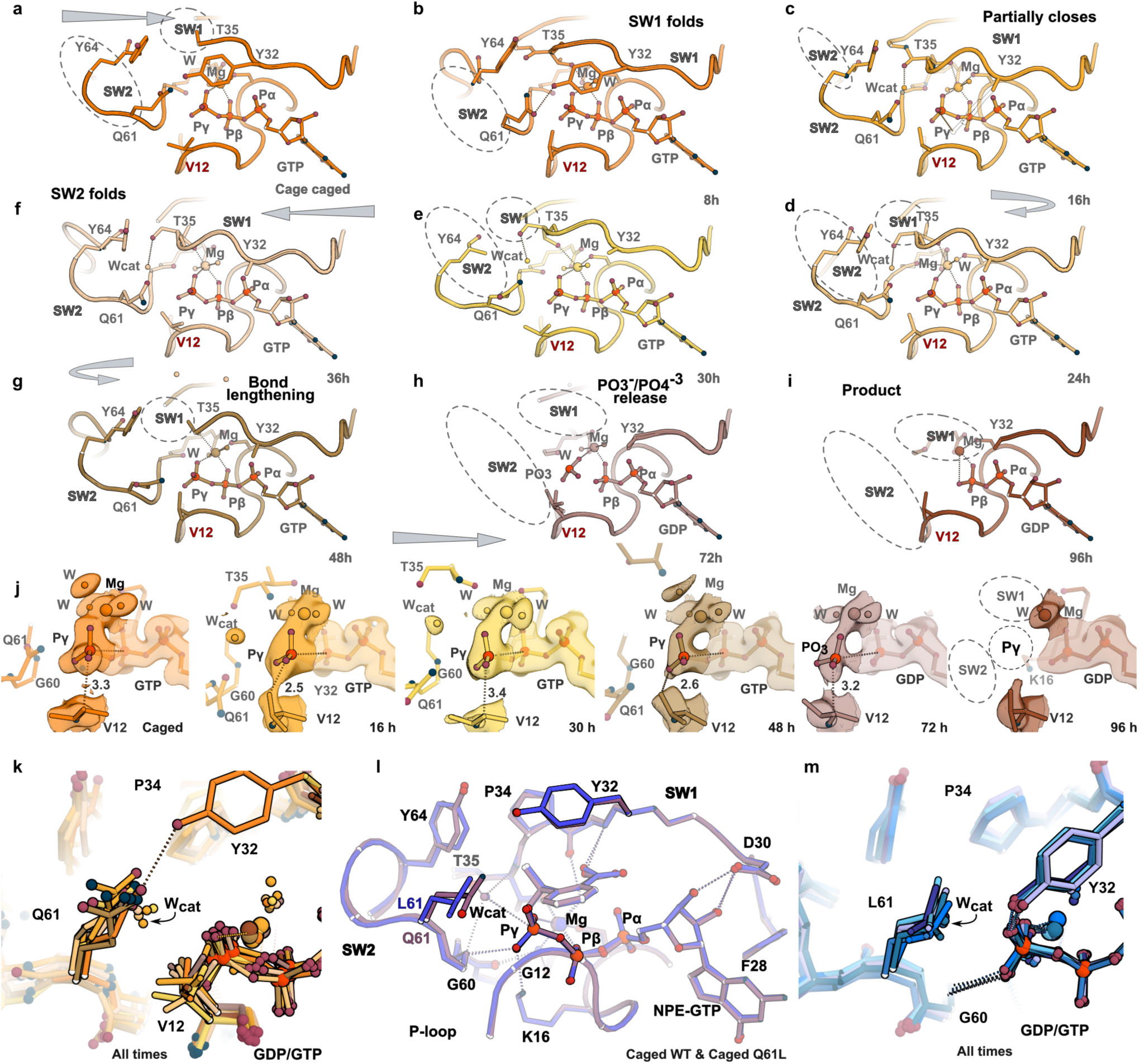
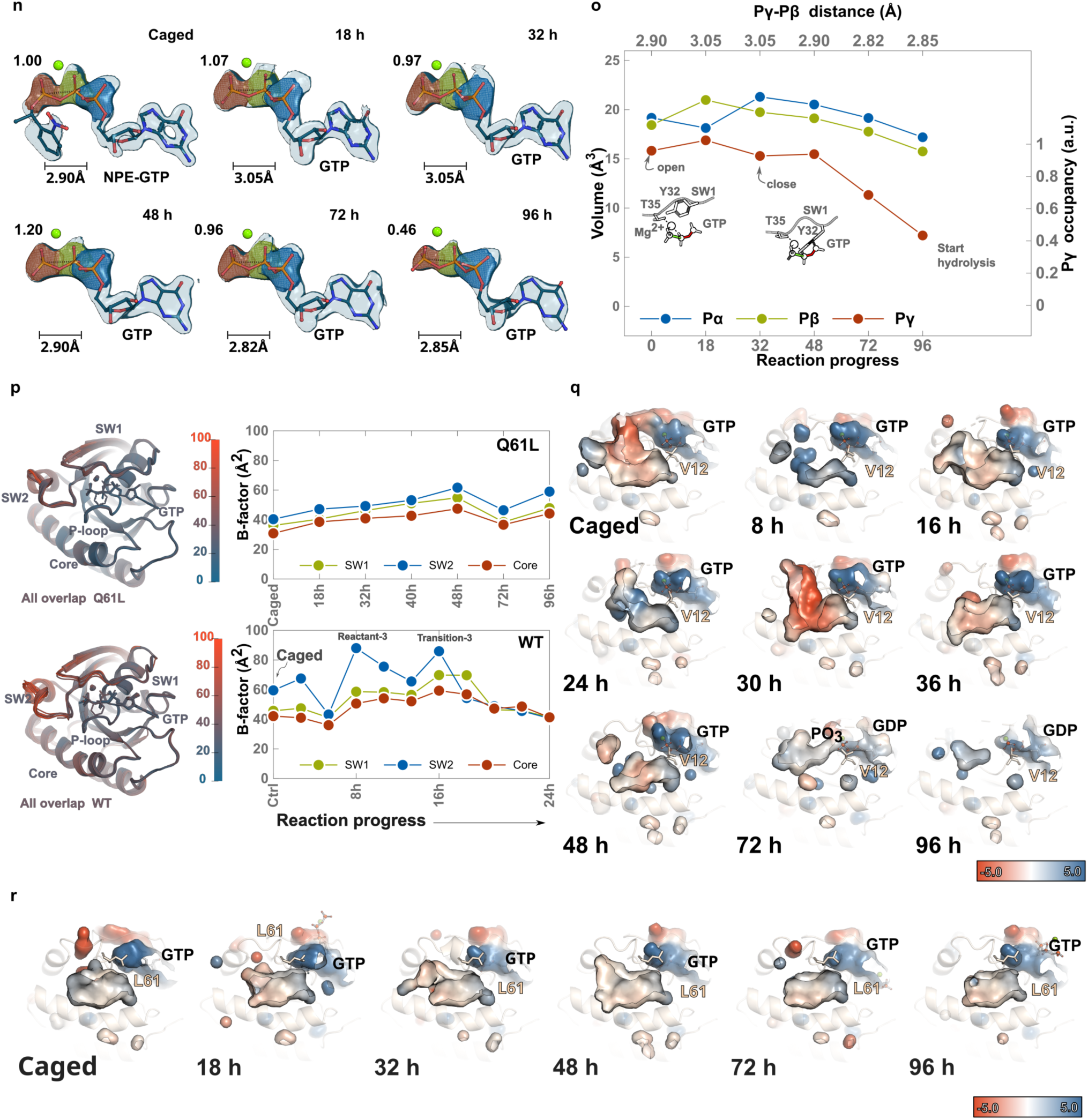
Structural basis for slow GTP hydrolysis observed in the G12V and Q61L mutants. **a**-**i**, Cartoon and ball and stick representation (*<u>switch view</u>*) of the ten states (from caged to 96-hours) during G12V GTP hydrolysis. Dashed ovals represent unfolded switch regions. **j**, Electron density 2Fo-Fc map contoured at 1.5 sigma for G12V γ-phosphate during the hydrolysis reaction. **k**, Illustration of positions adopted by Val^12^ and Gln^61^ during the hydrolysis reaction. **l**, Overlay between NPE-GTP bound Q61L (blue) and WT (purple). **m**, Illustration of positions adopted by Leu^61^ and Tyr^32^ for all time points of the reaction. Note the low mobility of the leucine residue and residues in the active site. **n**, Polder maps contoured at 4.5 0 omitting GTP from the calculation. Blue, green and red spheres correspond to α-β- and γ-phosphates, respectively. Bar illustrates the β-γ bond distance. **o**, Occupancy of the γ-phosphate as a function of the reaction process (**p**) B-factor comparison for Q61L (top panel) and WT (bottom panel) structural states. (**q,r**) Surface electrostatic representation (colored according to electrostatic potential (from −5 kT/e, red, to +5 kT/e, blue) of the intermediate state SW2 cryptic pockets illustrating morphology changes at different times during GTP hydrolysis for G12V (**q**) and Q61L (**r**)

**Extended Data Fig. 7:**
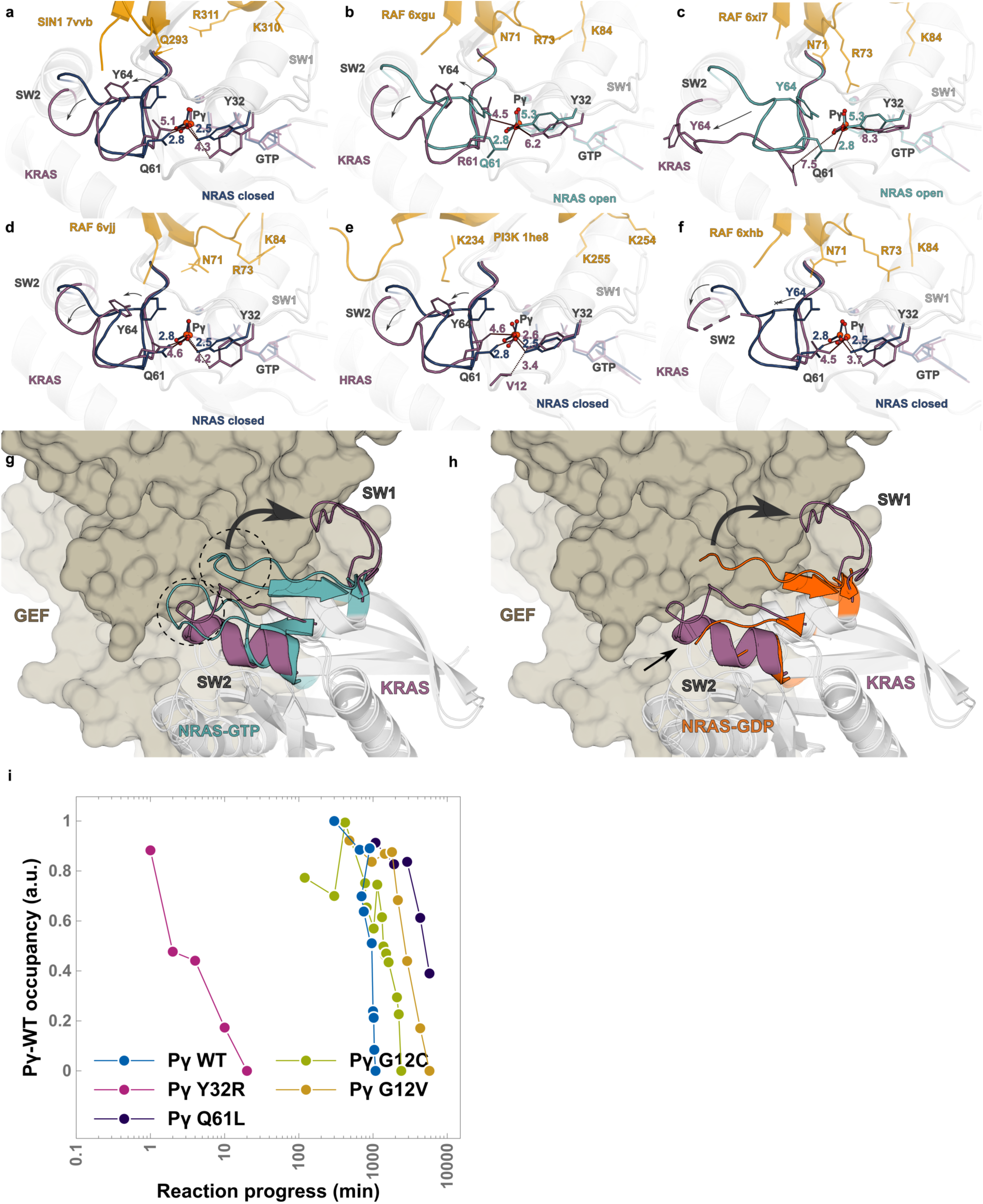
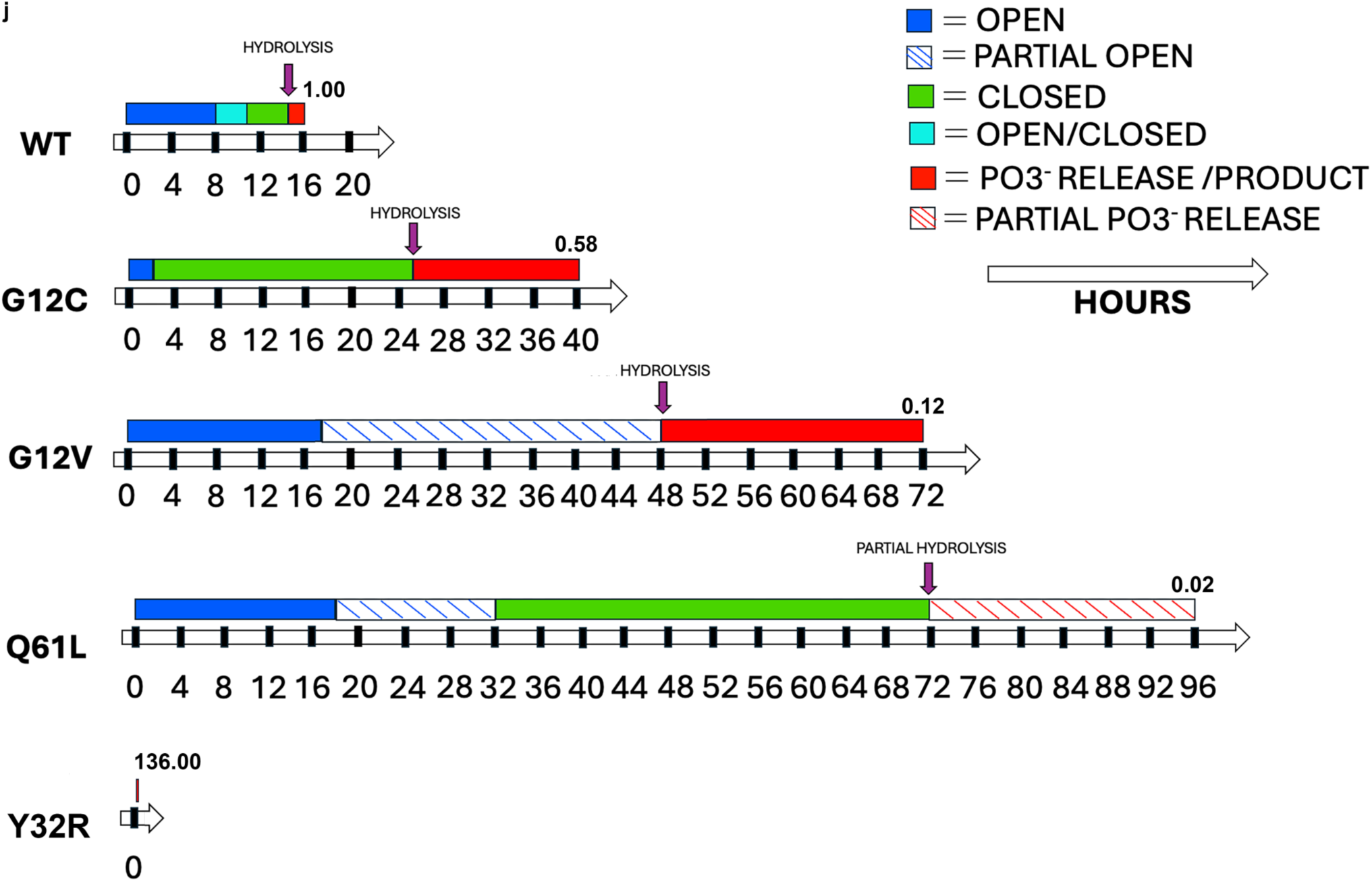
Effectors can bind open and closed states. **a**-**f**, Overlay of K-RAS-Effector complexes with WT N-RAS state VIII (blue) or state III (cyan) illustrating that effectors can potentially bind to both open and closed states. Name of each effector and PDB:ID coordinate files are indicated. Arrows indicate the motion of SW2 residues (open or closed conformations) upon effector binding. **g**,**h**, Cartoon and surface representation of the overlay between K-RAS-SOS complex (purple cartoon and light brown surface, respectively) with the N-RAS GTP complex (state III, cyan) and N-RAS GDP complex (state XIV, orange) illustrating potential clashes with folded switches in the GTP-bound structure (**g**), but minimal clashes (small arrow) when the switches are disordered in the GDP product state (h). The large arrow indicates motion of SW1 residues upon GEF binding. **i**, Log plot of the reaction progress time (hours) vs γ-phosphate occupancy for Y32R (pink), WT (light blue), G12C (green), G12V (yellow) and Q61L (dark blue). **j**, Schematic of the reaction progress during GTP hydrolysis for WT, G12C, G12V, Q61L and Y32R to illustrate the different duration of their intermediate states.

**Extended Data Fig. 8:**
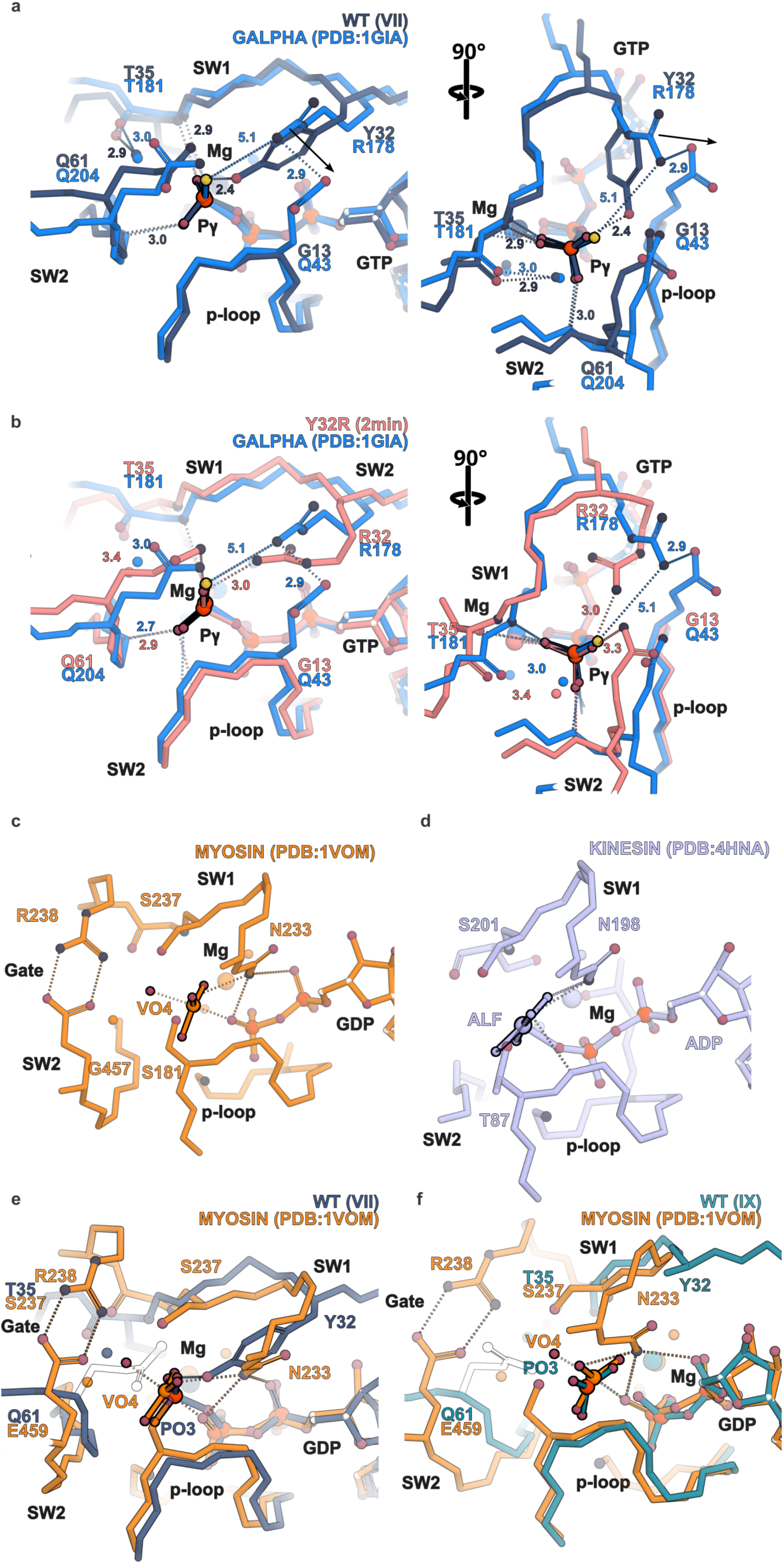
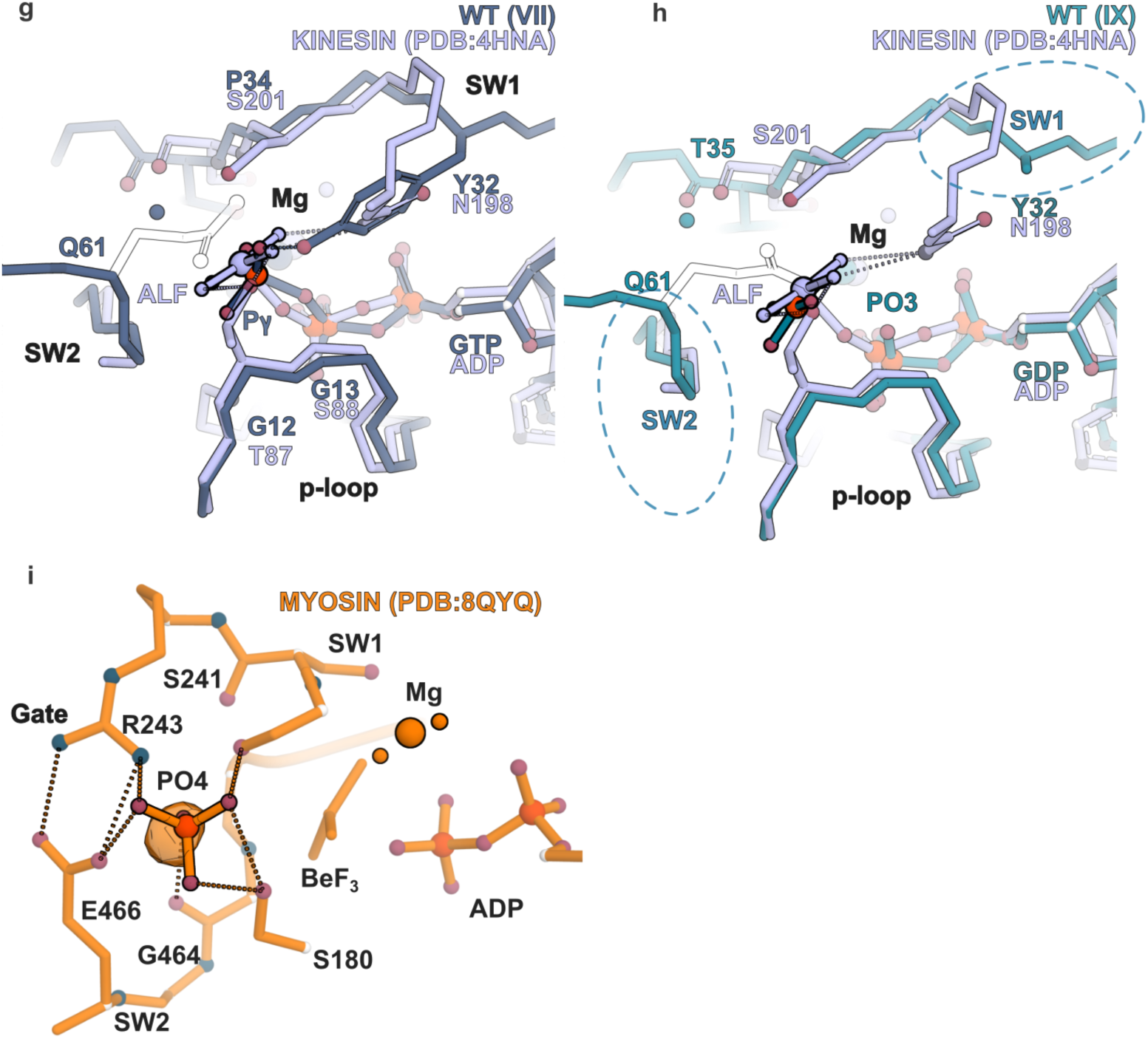
Conservation of the GTP and ATP hydrolysis mechanism. **a**, Two views, phosphate (left) and switch (right), of the overlay between the WT N-RAS structure (state VII, dark blue) and Gα_s_ (PDB:ID 1GIA) (light blue). SW1, SW2 and P loops architecture are well-conserved; Gα_s_, Arg^178^ replaces Tyr^32^ increasing its hydrolytic rate however, its distance is modulated by P-loop residue Glu^43^. **b**, Two views, *<u>phosphate</u>* (left) and *<u>switch</u>* (right), of the overlay between the Y32R mutant N-RAS structure (pink) and Gα_s_ (PDB:ID 1GIA) (light blue). Notice the shorter γ-phosphate distance in Y32R (3 Å vs 5.1 Å). **c**, Switch view of *Dictyostelium discoideum* vanadate-trapped (transition state structure) myosin I (PDB:ID 1VOM) illustrating gate residues R238 (SW1) and G457 (SW2) and the position of the catalytic N233 (SW1); VO4 is stabilized by multiple H-bonds with neighboring residues. Dashes indicate H-bonds (<3.5 Å). (d) Switch view of kinesin’s motor domain in complex with ADP and AlF4, illustrating AlF4 stabilizing interactions by P-loop and SW1 residues and conservation of the catalytic asparagine residue. Overlay of myosin PDB:ID VOM1 and NRAS state VII (**e**) and IX (**f**) structures illustrating conservation of the active sites including position of SW1 Y32 and (N233), P-loop and SW2 residues. Overlay of PDB:ID 4HNA and NRAS state VII (**g**) and IX (**h**) structures illustrating conservation of the active sites including position of SW1 residues Y32 and (N198). **i**, Structure of the motor domain of the pre-power stroke of β-cardiac myosin VII (PDB:ID 8QYQ) bound to BeF3. Orange surface indicates density (contoured at 20) for W_cat._ A modeled PO4^3-^ illustrates its potential position before release.

**Extended Data Table 1.** Activation free energy calculation. Computed changes of rates of GTP hydrolysis in NRAS mutants compared to WT NRAS.

| | $\Delta G^\ddagger$<br>(kcal/mol) | $\Delta\Delta G^\ddagger$ vs WT<br>(kcal/mol) | $K_{\text{hydr}}/K_{\text{hydr\_NRAS\_WT}}$ |
| --- | --- | --- | --- |
| NRAS WT <sup>close</sup> | 7.97 | 0 | 1 |
| NRAS WT <sup>open</sup> | 9.18 | 1.21 | 0.12 |
| NRAS Y32R | 5.15 | -2.82 | 135.68 |
| NRAS G12V | 9.21 | 1.24 | 0.115 |
| NRAS Q61L | 10.19 | 2.22 | 0.021 |
| NRAS G12C | 8.51 | 0.54 | 0.58 |
| SOLV | 18.92 | 10.95 | 5.24E-09 |

## References

1 Bourne, H. R., Sanders, D. A. & McCormick, F. The GTPase superfamily: a conserved switch for diverse cell functions. Nature 348, 125–132 (1990). 10.1038/348125a0

2 Langbeheim, H., Shih, T. Y. & Scolnick, E. M. Identification of a normal vertebrate cell protein related to the p21 src of Harvey murine sarcoma virus. Virology 106, 292–300 (1980). 10.1016/0042-6822(80)90252-4

3 Santos, E. et al. Malignant activation of a K-ras oncogene in lung carcinoma but not in normal tissue of the same patient. Science 223, 661–664 (1984). 10.1126/science.6695174

4 Der, C. J., Krontiris, T. G. & Cooper, G. M. Transforming genes of human bladder and lung carcinoma cell lines are homologous to the ras genes of Harvey and Kirsten sarcoma viruses. Proc Natl Acad Sci U S A 79, 3637–3640 (1982). 10.1073/pnas.79.11.3637

5 Gibbs, J. B., Sigal, I. S., Poe, M. & Scolnick, E. M. Intrinsic GTPase activity distinguishes normal and oncogenic ras p21 molecules. Proc Natl Acad Sci U S A 81, 5704–5708 (1984). 10.1073/pnas.81.18.5704

6 Trahey, M. & McCormick, F. A cytoplasmic protein stimulates normal N-ras p21 GTPase, but does not affect oncogenic mutants. Science 238, 542–545 (1987). 10.1126/science.2821624

7 Singhal, A., Li, B. T. & O’Reilly, E. M. Targeting KRAS in cancer. Nat Med 30, 969–983 (2024). 10.1038/s41591-024-02903-0

8 Neal, S. E., Eccleston, J. F. & Webb, M. R. Hydrolysis of Gtp by P21nras, the Nras Protooncogene Product, Is Accompanied by a Conformational Change in the Wild-Type Protein - Use of a Single Fluorescent-Probe at the Catalytic Site. P Natl Acad Sci USA 87, 3562–3565 (1990). DOI10.1073/pnas.87.9.3562

9 Baier, F. et al. Cryptic genetic variation shapes the adaptive evolutionary potential of enzymes. Elife 8 (2019). 10.7554/eLife.40789

10 Hoshino, M., Kawakita, M. & Hattori, S. Characterization of a Factor That Stimulates Hydrolysis of Gtp Bound to Ras Gene Product-P21 (Gtpase-Activating Protein) and Correlation of Its Activity to Cell-Density. Mol Cell Biol 8, 4169–4173 (1988). Doi 10.1128/Mcb.8.10.4169

11 Calixto, A. R., Moreira, C. & Kamerlin, S. C. L. Recent Advances in Understanding Biological GTP Hydrolysis through Molecular Simulation. Acs Omega 5, 4380–4385 (2020). 10.1021/acsomega.0c00240

12 Kotting, C., Kallenbach, A., Suveyzdis, Y., Wittinghofer, A. & Gerwert, K. The GAP arginine finger movement into the catalytic site of Ras increases the activation entropy. Proc Natl Acad Sci U S A 105, 6260–6265 (2008). 10.1073/pnas.0712095105

13 McCormick, F. et al. Interaction of ras p21 proteins with GTPase activating protein. Cold Spring Harb Symp Quant Biol 53 **Pt** **2**, 849–854 (1988). 10.1101/sqb.1988.053.01.097

14 Scheffzek, K. et al. The Ras-RasGAP complex: structural basis for GTPase activation and its loss in oncogenic Ras mutants. Science 277, 333–338 (1997). 10.1126/science.277.5324.333

15 Milburn, M. V. et al. Molecular switch for signal transduction: structural differences between active and inactive forms of protooncogenic ras proteins. Science 247, 939–945 (1990). 10.1126/science.2406906

16 Pai, E. F. et al. Refined crystal structure of the triphosphate conformation of H-ras p21 at 1.35 A resolution: implications for the mechanism of GTP hydrolysis. EMBO J 9, 2351–2359 (1990). 10.1002/j.1460-2075.1990.tb07409.x

17 Parker, M. I., Meyer, J. E., Golemis, E. A. & Dunbrack, R. L. Delineating the RAS Conformational Landscape. Cancer Res 82, 2485–2498 (2022). 10.1158/0008-5472.Can-22-0804

18 Maegley, K. A., Admiraal, S. J. & Herschlag, D. Ras-catalyzed hydrolysis of GTP: a new perspective from model studies. Proc Natl Acad Sci U S A 93, 8160–8166 (1996). 10.1073/pnas.93.16.8160

19 Li, G. & Zhang, X. C. GTP hydrolysis mechanism of Ras-like GTPases. J Mol Biol 340, 921–932 (2004). 10.1016/j.jmb.2004.06.007

20 Calixto, A. R. et al. GTP Hydrolysis Without an Active Site Base: A Unifying Mechanism for Ras and Related GTPases. J Am Chem Soc 141, 10684–10701 (2019). 10.1021/jacs.9b03193

21 Prakash, P. & Gorfe, A. A. Overview of simulation studies on the enzymatic activity and conformational dynamics of the GTPase Ras. Mol Simul 40, 839–847 (2014). 10.1080/08927022.2014.895000

22 Carvalho, A. T., Szeler, K., Vavitsas, K., Aqvist, J. & Kamerlin, S. C. Modeling the mechanisms of biological GTP hydrolysis. Arch Biochem Biophys 582, 80–90 (2015). 10.1016/j.abb.2015.02.027

23 Lin, J., Gerwert, K. & Kotting, C. A modified infrared spectrometer with high time resolution and its application for investigating fast conformational changes of the GTPase Ras. Appl Spectrosc 68, 531–535 (2014). 10.1366/13-07320

24 Rudack, T., Xia, F., Schlitter, J., Kötting, C. & Gerwert, K. Ras and GTPase-activating protein (GAP) drive GTP into a precatalytic state as revealed by combining FTIR and biomolecular simulations. P Natl Acad Sci USA 109, 15295–15300 (2012). 10.1073/pnas.1204333109

25 Tidyman, W. E. & Rauen, K. A. The RASopathies: developmental syndromes of Ras/MAPK pathway dysregulation. Curr Opin Genet Dev 19, 230–236 (2009). 10.1016/j.gde.2009.04.001

26 Duarte, F., Barrozo, A., Aqvist, J., Williams, N. H. & Kamerlin, S. C. The Competing Mechanisms of Phosphate Monoester Dianion Hydrolysis. J Am Chem Soc 138, 10664–10673 (2016). 10.1021/jacs.6b06277

27 Barrozo, A. et al. Computer simulations of the catalytic mechanism of wild-type and mutant beta-phosphoglucomutase. Org Biomol Chem 16, 2060–2073 (2018). 10.1039/c8ob00312b

28 Scheidig, A. J. et al. X-ray crystal structure analysis of the catalytic domain of the oncogene product p21H-ras complexed with caged GTP and mant dGppNHp. J Mol Biol 253, 132–150 (1995). 10.1006/jmbi.1995.0541

29 Amyes, T. L. & Richard, J. P. Specificity in transition state binding: the Pauling model revisited. Biochemistry 52, 2021–2035 (2013). 10.1021/bi301491r

30 Araki, M. et al. Solution structure of the state 1 conformer of GTP-bound H-Ras protein and distinct dynamic properties between the state 1 and state 2 conformers. J Biol Chem 286, 39644–39653 (2011). 10.1074/jbc.M111.227074

31 Moghadamchargari, Z. et al. Intrinsic GTPase Activity of K-RAS Monitored by Native Mass Spectrometry. Biochemistry 58, 3396–3405 (2019). 10.1021/acs.biochem.9b00532

32 Ahmadian, M. R. et al. Guanosine triphosphatase stimulation of oncogenic Ras mutants. Proc Natl Acad Sci U S A 96, 7065–7070 (1999). 10.1073/pnas.96.12.7065

33 Rudack, T., Xia, F., Schlitter, J., Kotting, C. & Gerwert, K. The role of magnesium for geometry and charge in GTP hydrolysis, revealed by quantum mechanics/molecular mechanics simulations. Biophys J 103, 293–302 (2012). 10.1016/j.bpj.2012.06.015

34 Okimoto, N. et al. Theoretical studies of the ATP hydrolysis mechanism of myosin. Biophys J 81, 2786–2794 (2001). 10.1016/S0006-3495(01)75921-8

35 Lin, G. et al. Structural basis of transcription: RNA polymerase II substrate binding and metal coordination using a free-electron laser. Proc Natl Acad Sci U S A 121, e2318527121 (2024). 10.1073/pnas.2318527121

36 Vergara, S. et al. Structural basis of deoxynucleotide addition by HIV-1 RT during reverse transcription. Nat Commun 15, 10553 (2024). 10.1038/s41467-024-54618-y

37 DeFeo, D. et al. Analysis of two divergent rat genomic clones homologous to the transforming gene of Harvey murine sarcoma virus. Proc Natl Acad Sci U S A 78, 3328–3332 (1981). 10.1073/pnas.78.6.3328

38 Shimizu, K. et al. Three human transforming genes are related to the viral ras oncogenes. Proc Natl Acad Sci U S A 80, 2112–2116 (1983). 10.1073/pnas.80.8.2112

39 Der, C. J., Finkel, T. & Cooper, G. M. Biological and biochemical properties of human rasH genes mutated at codon 61. Cell 44, 167–176 (1986). 10.1016/0092-8674(86)90495-2

40 Hunter, J. C. et al. Biochemical and Structural Analysis of Common Cancer-Associated KRAS Mutations. Mol Cancer Res 13, 1325–1335 (2015). 10.1158/1541-7786.MCR-15-0203

41 Richard, J. P., Amyes, T. L., Goryanova, B. & Zhai, X. Enzyme architecture: on the importance of being in a protein cage. Curr Opin Chem Biol 21, 1–10 (2014). 10.1016/j.cbpa.2014.03.001

42 Tran, T. H. et al. KRAS interaction with RAF1 RAS-binding domain and cysteine-rich domain provides insights into RAS-mediated RAF activation. Nat Commun 12, 1176 (2021). 10.1038/s41467-021-21422-x

43 Pacold, M. E. et al. Crystal structure and functional analysis of Ras binding to its effector phosphoinositide 3-kinase gamma. Cell 103, 931–943 (2000). 10.1016/s0092-8674(00)00196-3

44 Zheng, Y. et al. Structural insights into Ras regulation by SIN1. Proc Natl Acad Sci U S A 119, e2119990119 (2022). 10.1073/pnas.2119990119

45 Kiani, F. A. & Fischer, S. Stabilization of the ADP/metaphosphate intermediate during ATP hydrolysis in pre-power stroke myosin: quantitative anatomy of an enzyme. J Biol Chem 288, 35569–35580 (2013). 10.1074/jbc.M113.500298

46 Santos, E. & Nebreda, A. R. Structural and functional properties of ras proteins. FASEB J 3, 2151–2163 (1989). 10.1096/fasebj.3.10.2666231

47 Woods, A. S. & Ferre, S. Amazing stability of the arginine-phosphate electrostatic interaction. J Proteome Res 4, 1397–1402 (2005). 10.1021/pr050077s

48 Stephen, A. G., Esposito, D., Bagni, R. K. & McCormick, F. Dragging ras back in the ring. Cancer Cell 25, 272–281 (2014). 10.1016/j.ccr.2014.02.017

49 Kanade, M., Chakraborty, S., Shelke, S. S. & Gayathri, P. A Distinct Motif in a Prokaryotic Small Ras-Like GTPase Highlights Unifying Features of Walker B Motifs in P-Loop NTPases. J Mol Biol 432, 5544–5564 (2020). 10.1016/j.jmb.2020.07.024

50 Thomas, C. J. et al. Uncoupling conformational change from GTP hydrolysis in a heterotrimeric G protein alpha-subunit. Proc Natl Acad Sci U S A 101, 7560–7565 (2004). 10.1073/pnas.0304091101

51 Cross, R. A. Review: Mechanochemistry of the kinesin-1 ATPase. Biopolymers 105, 476–482 (2016). 10.1002/bip.22862

52. Mueller, M. P. & Goody, R. S. Review: Ras GTPases and myosin: Qualitative conservation and quantitative diversification in signal and energy transduction. Biopolymers 105, 422-430 (2016). 10.1002/bip.22840

53 Ge, J., Huang, F. & Nesmelov, Y. E. Metal cation controls phosphate release in the myosin ATPase. Protein Sci 26, 2181–2186 (2017). 10.1002/pro.3267

54 Fisher, A. J. et al. X-ray structures of the myosin motor domain of Dictyostelium discoideum complexed with MgADP.BeFx and MgADP.AlF4. Biochemistry 34, 8960–8972 (1995). 10.1021/bi00028a004

55 Gigant, B. et al. Structure of a kinesin-tubulin complex and implications for kinesin motility. Nat Struct Mol Biol 20, 1001–1007 (2013). 10.1038/nsmb.2624

56 Moretto, L. et al. Multistep orthophosphate release tunes actomyosin energy transduction. Nat Commun 13, 4575 (2022). 10.1038/s41467-022-32110-9

57 Vagin, A. & Teplyakov, A. Molecular replacement with MOLREP. Acta Crystallogr D Biol Crystallogr 66, 22–25 (2010). 10.1107/S0907444909042589

58 Emsley, P. & Cowtan, K. Coot: model-building tools for molecular graphics. Acta Crystallogr D Biol Crystallogr 60, 2126–2132 (2004). 10.1107/S0907444904019158

59 Liebschner, D. et al. Macromolecular structure determination using X-rays, neutrons and electrons: recent developments in Phenix. Acta Crystallogr D Struct Biol 75, 861–877 (2019). 10.1107/S2059798319011471

60 Emsley, P., Lohkamp, B., Scott, W. G. & Cowtan, K. Features and development of Coot. Acta Crystallogr D Biol Crystallogr 66, 486–501 (2010). 10.1107/S0907444910007493

61 Meng, E. C. et al. UCSF ChimeraX: Tools for structure building and analysis. Protein Sci 32, e4792 (2023). 10.1002/pro.4792

62 Goddard, T. D. et al. UCSF ChimeraX: Meeting modern challenges in visualization and analysis. Protein Sci 27, 14–25 (2018). 10.1002/pro.3235

63 Moffat, J. et al. A lentiviral RNAi library for human and mouse genes applied to an arrayed viral high-content screen. Cell 124, 1283–1298 (2006). 10.1016/j.cell.2006.01.040

64 Yochum, Z. A. et al. A First-in-Class TWIST1 Inhibitor with Activity in Oncogene-Driven Lung Cancer. Mol Cancer Res 15, 1764–1776 (2017). 10.1158/1541-7786.MCR-17-0298

65 Yochum, Z. A. et al. Targeting the EMT transcription factor TWIST1 overcomes resistance to EGFR inhibitors in EGFR-mutant non-small-cell lung cancer. Oncogene 38, 656–670 (2019). 10.1038/s41388-018-0482-y

66 Laskowski, R. A. & Swindells, M. B. LigPlot+: multiple ligand-protein interaction diagrams for drug discovery. J Chem Inf Model 51, 2778–2786 (2011). 10.1021/ci200227u

67 Wallace, A. C., Laskowski, R. A. & Thornton, J. M. LIGPLOT: a program to generate schematic diagrams of protein-ligand interactions. Protein Eng 8, 127–134 (1995). 10.1093/protein/8.2.127

68 Kim, D. et al. Pan-KRAS inhibitor disables oncogenic signalling and tumour growth. Nature 619, 160–166 (2023). 10.1038/s41586-023-06123-3

